# Transient interdomain interactions modulate the monomeric structural ensemble and self-assembly of Huntingtin Exon 1

**DOI:** 10.1101/2024.05.03.592468

**Authors:** Priyesh Mohanty, Tien Minh Phan, Jeetain Mittal

## Abstract

Polyglutamine expansion (≥ 36 residues) within the N-terminal exon-1 of Huntingtin (Httex1) leads to Huntington’s disease, a neurodegenerative condition marked by the presence of intranuclear Htt inclusions. Notably, the polyglutamine tract in Httex1 is flanked by an N-terminal coiled-coil domain - N17 (17 amino acids), which undergoes self-association to promote the formation of soluble Httex1 oligomers and brings the aggregation-prone polyQ tracts in close spatial proximity. However, the mechanisms underlying the subsequent conversion of soluble oligomers into insoluble β-rich aggregates with increasing polyQ length, remain unclear. Current knowledge suggests that expansion of the polyQ tract increases its helicity, and this favors its oligomerization and aggregation. In addition, studies utilizing photocrosslinking, conformation-specific antibodies and a stable coiled-coil heterotetrametric system fused to polyQ indicate that domain “cross-talk” (i.e., interdomain interactions) may play a role in the emergence of toxic conformations and the conversion of Httex1 oligomers into fibrillar aggregates. Here, we performed extensive atomistic molecular dynamics (MD) simulations (aggregate time ∼ 0.7 ms) to uncover the interplay between structural transformation and domain “cross-talk” on the conformational ensemble and oligomerization landscape of Httex1. Notably, our MD-derived ensembles of N17-polyQ monomers validated against ^13^C NMR chemical shifts indicated that in addition to elevated α- helicity, polyQ expansion also favors transient, interdomain (N17-polyQ) interactions which result in the emergence of β-sheet conformations. Further, interdomain interactions competed with increased polyQ tract α-helicity to modulate the stability of N17-mediated dimers and thereby promoted a heterogenous dimerization landscape. Finally, we observed that the C-terminal proline-rich domain (PRD) promoted condensation of Httex1 through self-interactions involving its P_10_/P_11_ tracts while also interacting with N17 to suppress its α-helicity. In summary, our study demonstrates a significant role for domain “cross-talk” in modulating the monomeric structural ensemble and self-assembly of Httex1.

## Introduction

There are nine polyglutamine (polyQ)-associated neurodegenerative disorders which are known to arise as a result of CAG trinucleotide repeat expansion within the polyglutamine-encoding tract of a specific gene.^1–3^ Huntington’s disease (HD) is a polyglutamine-related disorder marked by the presence intranuclear inclusions (aggregates) of the protein - Huntingtin (Htt) in striatal neurons.^4,5^ At present, there is no cure available for HD.^6,7^ Huntingtin (Htt) is a 348 kDa multifunctional protein which shuttles between nucleus and cytoplasm, is broadly involved in the development of the nervous system, brain-derived neutrophic factor (BDNF) production and transport, and cell adhesion.^8^ The loss of native Htt function and its toxic gain-of-function via length expansion and aggregation of the polyglutamine tract (>36 residues) within the N-terminal Htt exon-1 (Httex1) is associated with HD.^9,10^ The length of polyQ expansion inversely correlates with the age of disease onset.^11^ Notably, Httex1 fragments generated through proteolytic cleavage are observed in intranuclear inclusions^12^ and *in vivo* expression of mutant Httex1 was found to reproduce the key features of HD in mice models.^13,14^ Not surprisingly, the structural and conformational changes associated with polyglutamine expansion^15–17^ and the precise role of flanking regions in Httex1 oligomerization^18^, fibrillar aggregation^19^ and cellular toxicity^20^ has been the subject of intense debate and investigations.^21,22^

Httex1 is an intrinsically disordered polypeptide (91 amino acids) comprising of (i) an N- terminal α-helical domain (N17: 17 amino acids), a central polyglutamine (polyQ_n_) tract which is aggregation-prone and a C-terminal proline rich domain (PRD: 51 amino acids) **(Fig. 1A)**. The polyglutamine flanking domains (N17/PRD) exert opposing effects on fibrillar aggregation; N17 promotes fibrillation and suppresses the formation of non-fibrillar species, while PRD generally disfavors aggregation and promotes the formation of soluble oligomers.^19,23–25^ In terms of the classical nucleation model for polyglutamine amyloid formation,^26^ Wetzel and colleagues determined that the size of the critical nucleus required to template the formation of β-rich aggregates of dilysine-flanked polyglutamine peptides *in vitro* depends upon the length of polyQ: the nuclei for >Q_25_ are monomeric while the smaller fragments are observed to form larger nuclei (n=2-4).^27^ These findings were further corroborated in a computational study by Wolynes and co-workers based on simulated free energy landscapes of polyglutamine aggregation computed using a coarse-grained protein model.^28^ Computational modeling also suggests that the nucleation of smaller polyQ fragments promotes elongation and fibrillation by reducing the conformational entropy of interacting monomers in a concentration-dependent manner.^29^ Recently, Halfmann and colleagues confirmed the formation of a monomeric nucleus *in vivo* for the pathogenic polyQ_60_ fragment and determined its structural features using an integrative modeling approach.^30^

**Figure 1.**
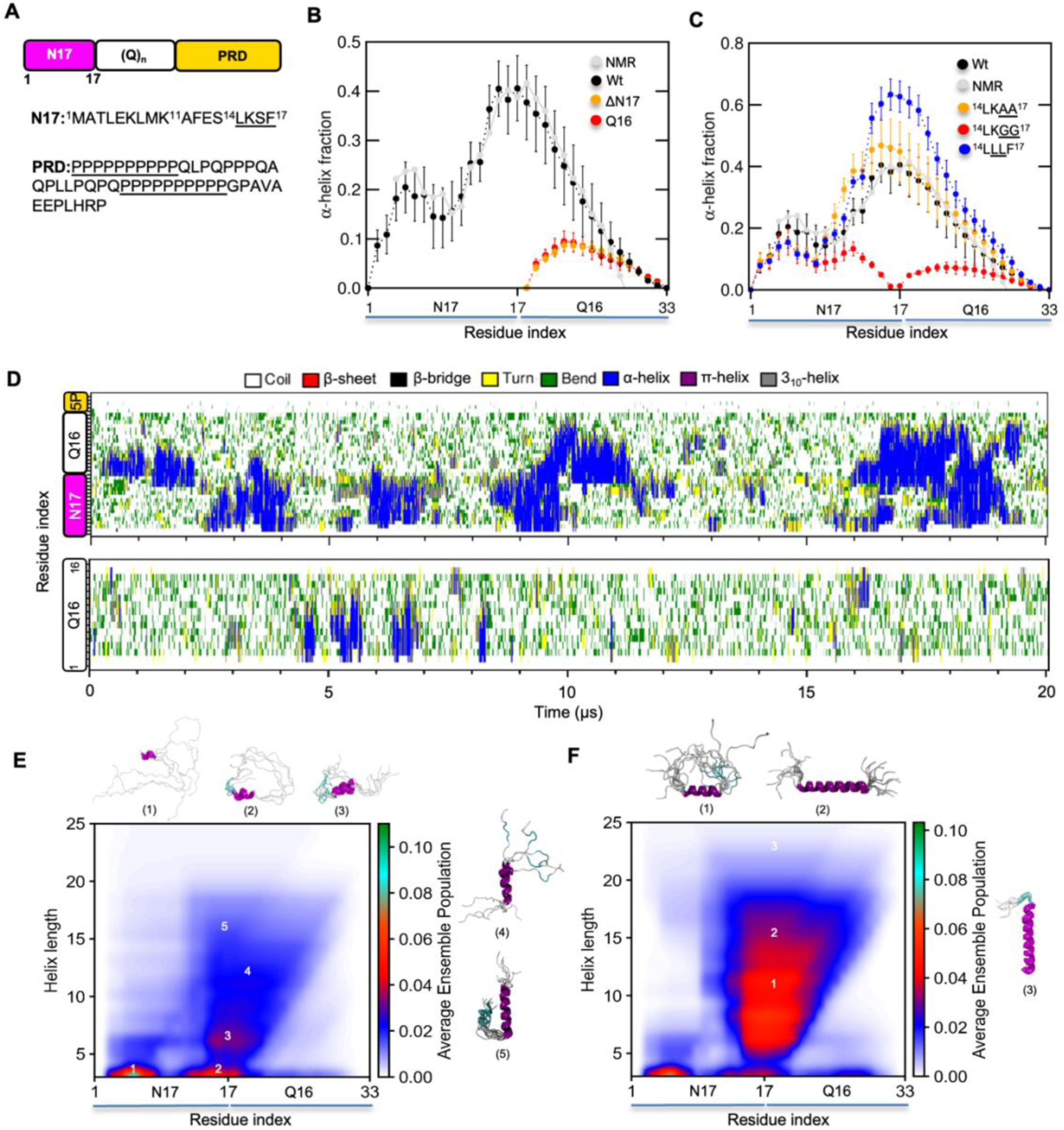
Analysis of N17-Q_16_ conformational ensemble illustrates the structural cooperativity between N17 and polyQ regions. **A.** Schematic showing the organization of Httex1 into three regions - N17, polyQ and PRD (top), and the corresponding sequences of N17/PRD domains (bottom). The length of polyQ region in the wild-type Huntingtin protein is ∼23 residues. **B.** Comparison of per-residue α-helix fractions for N17-Q_16_ wild-type and its deletion constructs with respect to NMR. Helical fractions were computed using the DSSP algorithm and compared to SSP scores for the helical propensity (grey) computed from experimental ^13^C chemical shifts. Error bars denote std. error of mean calculated over three independent trajectories. **C.** Same as in panel B for N17-Q_16_ wild-type and ^14^LKSF^17^ mutants. **D.** Secondary structure variation as a function of time in a representative N17-Q_16_ (top) and Q16 (bottom) trajectory. **E.** SS-map of N17-Q16 wild-type computed from an aggregate trajectory (∼63 μs) indicating the probability of various helical length across N17 and polyQ regions. Representative α-helical conformations corresponding to numbered regions of the SS-map are shown in cartoon representation. **F.** Same as in E for ^14^LLLF^17^ mutant.

N17 significantly enhances the rate of polyQ fibrillation *in vitro and in vivo* through a complex, multistep mechanism involving small (low molecular weight) prenucleation oligomers which undergo further rearrangement to promote amyloid nucleation.^31,32^ A kinetically competing, N17-independent aggregation pathway also exists for Huntingtin exon 1 which resembles that of simple polyglutamine and dominates upon disruption of N17 association or its partial excision.^33^ Notably, solid-state NMR experiments concluded that the core of Httex1 fibrils was structurally similar to simple polyglutamine fragments, comprising of an inter-digitating hydrogen bond network formed between adjacent layers of polyQ β-hairpins through glutamine sidechains while the flanking N17 remains helical.^34–36^ While higher-order oligomerization through N17 significantly increases the local concentration of polyQ tracts (‘proximity’ model), such a mechanism alone fails to explain the enhanced rate of polyQ aggregation as forcing a high local concentration of polyQ by fusing it to homo-oligomeric partners suppresses its fibrillar aggregation *in vitro*^37^ and *in vivo.*^30^ Two additional models were proposed to explain the conversion of Httex1 oligomers into aggregates via polyQ expansion^37^: (i) ‘transformation’ model, i.e., the propagation of N17 helical secondary structure into the polyQ tract and, (ii) ‘domain cross-talk’ model, i.e., tertiary interdomain interactions between N17 and polyQ domains. A model of Httex1 amyloid aggregation based on photocross-linking experiments proposes that long-range N17-polyQ interactions result in a conformational switch which promotes rearrangement of the initial N17- mediated oligomers to form aggregation-competent nuclei.^32^

The relevance of interdomain interactions for Httex1 aggregation is further supported by *in vivo* experiments where the selective binding of a polyQ β-hairpin antibody to mutant Httex1 oligomers^38^ was favorable when atleast one of the flanking regions (N17 or PRD) were present.^23^ Nevertheless, ^42^the substantially faster aggregation of longer Httex1 constructs (>Q_7_) renders them less amenable to high resolution NMR experiments for studying conformational exchange and detecting interdomain interactions occurring with oligomeric assemblies.^39,40^ Alternatively, molecular dynamics (MD) simulations have the potential to complement solution experiments and provide a high-resolution picture of the conformational landscape and underlying mechanisms which facilitate the conversion of α-helical oligomers into amyloid aggregates. Towards this goal, numerous studies have attempted to investigate the structural and conformational changes in Httex1 monomers upon polyQ expansion using atomistic MD simulations.^41^ Nevertheless, these studies report conflicting results regarding the nature of pathogenic conformations due to: (i) inherent imbalances in secondary structure propensities (i.e., relative stabilities of α-helical versus β-conformations) for commonly-used protein force fields^42–44^ and, (ii) the tendency to generate excessively collapsed ensembles for unfolded and disordered proteins when simulated in implicit or primitive (three-site) explicit water models.^45,46,50–53^

In this study, we performed extensive MD simulations (aggregate time ∼ 0.7 ms) to evaluate the role of interdomain interactions on the monomeric structural ensemble, oligomerization, and condensation of Httex1. Using a state-of-the-art protein force field,^47^ our simulations yielded conformational ensembles in good agreement with residue-specific helical populations inferred from recent NMR experiments which employed site-specific isotope labeling of polyQ residues.^48,49^ With increasing polyQ length, we observed longer α-helices extending further into the polyglutamine tract along with the emergence of transient β-sheet conformations (<2% total population). Importantly, stable β-sheet conformations were not observed for a Q_46_- only fragment on a comparable simulation timescale, despite its low propensity for α-helix formation. These findings imply that interactions between N17 and polyQ domains are directly implicated in the enhancing the formation of β-sheet conformations which could act as precursors in the folding pathway enroute to the nucleus. Next, we investigated the interplay between helical transformation (i.e., elevated polyQ helicity) and interdomain interactions on the dimerization landscape of Httex1. We observed that an increase in polyQ length (7 to 16) allowed transient, intra and intermolecular N17-polyQ interactions to effectively outcompete the stabilizing effect of α-helical transformation and promote a heterogenous dimer ensemble. Finally, we observed that while the C-terminal proline-rich domain (PRD) does not alter the intrinsic α-helical propensity of Httex1 monomer (cis), it promotes condensation^50–52^ through intermolecular interactions involving P_10_/P_11_ tracts while also interacting with N17 (trans) to suppress its α-helicity. In conclusion, our results support a model wherein inter-domain interactions can exert opposing effects on Httex1 aggregation by both promoting the formation of aggregation-competent nuclei^32^ and condensation to limit aberrant phase transitions.^53,54^

## Results

### Validation of N17-Q_16_ simulation ensembles against NMR chemical shifts

The α-helical structure of N17 is associated with enhanced oligomerization and aggregation of Httex1 compared to their simple polyQ counterparts. Recent structural studies employing NMR spectroscopy with site-specific isotope labeling resolved the residue-level α-helical propensities of amino acids within the N17 and polyQ regions in Httex1 constructs with increasing polyQ length below (Q_16_) and above the pathogenic threshold (Q_46_).^55,56^ Such high resolution, residue-specific information serves an invaluable reference data set for the structural validation of Httex1 ensembles generated by MD force fields. We chose a balanced protein force field - AMBER03ws,^47^ to generate conformational ensembles of N17-(Q)_n_ monomers and dimers using multi-microsecond and enhanced sampling simulations (**Methods, Table S1**). Previous studies by us and others have firmly established the ability of AMBER03ws to accurately describe the global dimensions and local structural features of a wide variety of intrinsically disordered polypeptides (IDPs).^57,58^

We first performed multi-microsecond simulations of the N17-Q_16_ fragment with a minimal C-terminal stretch of five proline residues (P_5_). The mean probability distribution for the radius of gyration across three independent trajectories exhibited low standard errors, implying convergence of the N17-Q16 ensembles in terms of their global dimensions **(Fig. S1A)**. To validate the structural accuracy of the N17-Q_16_ ensemble, we first compared it with per-residue α-helical propensities computed from ^13^C chemical shifts using the SSP algorithm.^59^ Overall, mean per-residue helical fractions of N17-polyQ_16_ were found to be in excellent agreement with NMR across both N17 and Q_16_ regions **(Fig. 1B)**. Further, the direct comparison of ^13^C_α/β_ chemical shifts predicted using the SPARTA+ program (See Materials and Methods) indicated that the root mean square deviation (RMSD) over all residues with respect to experiment are within the prediction error **(Fig. S1B)**. The comparison of per-residue ^13^C secondary chemical shift difference (SCSD) with NMR shows excellent quantitative agreement (=<0.5 ppm) for several residues in the N17 region **(Fig S1C)**. Further, we observed a gradual decay in polyQ SCSD from the start (Q18) to the end position (Q33), a trend which is qualitatively consistent with experiment. As negative controls, we also simulated Q_16_-P_5_ and Q_16_ fragments; both exhibited substantially lower α-helicity compared to N17-polyQ_16_ which is consistent with experiment **(Fig. 1B)**.

NMR experiments indicated that a four residue motif in N17 (^14^LSKF^17^) structurally connects N17 and the downstream polyQ residues.^55^ Notably, these experiments revealed that ^14^LKSF^17^ uniquely favored the formation of bifurcated hydrogen bonds involving glutamine sidechains (i→i-4) in the adjacent polyQ region (aa:18-24) which could contribute to the elevated α-helical propensity.^48^ Similarly, an L_4_ motif flanking the Androgen receptor polyQ tract was found to promote bifurcated hydrogen bonds which stabilized its α-helical structure.^60^ Based on site-specific labeling of a few glutamine residues, it was observed that substitution of the wild-type motif - ^14^LKSF^17^ to either ^14^LKAA^17^ or ^14^LLLF^17^ enhanced helical stability while ^14^LKGG^17^ substitution significantly reduced the helicity of the polyQ region. We simulated all three ^14^LKSF^17^ variants to generate the per-residue helicity profiles of all mutants and assessed whether our force field could correctly capture the expected trends in their α-helicity profiles. The helix-promoting mutants showed an increase in α-helicity **(Fig. 1C)** starting from the C-terminal region of N17 (aa:11-17) leading into the polyQ region (aa:18-23). In contrast, ^14^LKGG^17^ disrupted the structural connectivity between N17 and polyQ regions, resulting in a complete loss of α-helical structure in the Q_16_ tract **(Fig. 1C)** and a corresponding increase in the population of coil conformations **(Fig. S1D)**. We also analyzed our ensembles for the prevalence of bifurcated hydrogen bonds in the polyQ tract among α-helical conformers (helix fraction >30% of total residues). While the overall occupancy of these hydrogen bonds was low, the observed trend correlated with their residue- level α-helical fractions: ^14^LLLF^17^ variant showed the higher hydrogen bond occupancies (>5%) in the central region (Q18/19) compared to ^14^LKAA^17^ and the wild-type sequence - ^14^LKSF^17^ **(Figure S1E)**.

Overall, the above comparison of N17-Q_16_ ensembles with high-resolution solution NMR data firmly establishes the suitability of our chosen model to generate structurally accurate ensembles of N17-polyQ fragments, assess the effect of point mutations on α-helix stability and analyze the mechanisms of helix initiation and propagation, as discussed in following section.

### Mechanism of α-helix nucleation and propagation into the polyQ tract

The high structural accuracy of our N17-Q_16_ ensemble motivated us to analyze the mechanisms of helix nucleation and propagation into the Q_16_ region which accounts for the observed residue- level α-helical populations. Accordingly, we first visualized the variation in secondary structure across both domains over the time course of the three independent trajectories (**Fig. 1D, S2A**). Upon careful inspection, we observed that α-helices generally appeared to nucleate in the region spanning the C-terminal N17 region (aa:11-17) and the adjacent Q_16_ tract (aa:18-23) and could further propagate in a bidirectional manner to form longer helices which persisted for several microseconds. However, these helices were less stable and did not persist for more than a microsecond. The N-terminal region of N17 (aa:3-10) formed transient helical structures (<1 μs) which were weakly associated with the formation of longer helices, suggestive of its structural decoupling from the downstream N17-polyQ sequence. Occasionally, N-terminal N17 also appeared to act as a nucleation site while also forming short α-helices which partially extended into the C-terminal N17 region **(Fig. S2A, bottom panel)**. In both N17-Q_16_ and Q_16_ trajectories, short α-helices (<8 residues) appeared within different stretches of polyQ sequence and usually persisted for less than 0.2 μs. In the case of Q_16_ trajectories, longer α-helices were also observed, although these typically did not persist beyond 1 μs, highlighting the critical requirement of N17 domain for stable α-helix nucleation and propagation into the polyQ tract (**Fig. 1D, S2B**).

The observations from time course analysis of N17-Q_16_ are summarized in the form of a α-helical secondary structure (SS) map^61^ which describes the sequence-level cooperativity underlying α-helix formation by reporting on the helical populations for a range of possible lengths. The SS map reveals that α-helical populations were the highest for single turns (<5 residues) in either N- or C-terminal region of N17, followed by a short α-helix in the central region (aa:11-23) which can propagate into further into the polyQ tract to form longer helices (**Fig. 1E**). In comparison, the ^14^LLLF^17^ variant showed a clear increase in the stability of longer α-helices (upto 15 residues) initiated from the central region as indicated their higher populations in SS map **(Fig. 1F).** The structural decoupling of the N-terminal N17 region was evident in N17-Q_16_ wild-type based on its reduced prevalence in longer helices (>8 residues) compared to the downstream N17-polyQ sequence and even more pronounced in case of the ^14^LLLF^17^ variant. Overall, the time course analysis of secondary structure formation and SS maps clearly establish a mechanism of helix initiation which is favored from the central region (aa:11-23) and its propagation into the polyQ tract leads to longer helices while the N-terminal N17 region (aa:3-10) has a low α-helix propensity and can be structurally uncoupled from the downstream sequence.

More recently, based on extensive evaluation and modification, Robustelli et al.^58^ proposed that the AMBER99SB-disp forcefield may represent a suitable force field choice for modeling IDP ensembles. Hence, we also simulated the N17-Q16 construct with AMBER99SB- disp and assessed its ability to accurately reproduce the NMR-derived helical fractions (Fig. S2). As evident from the per-residue α-helical fractions (**Fig. S3A**), the AMBER99SB-disp ensemble exhibits two major issues compared to NMR: (i) it incorrectly predicts the position for peak α- helicity and, (ii) overestimates the ɑ-helical fraction (by ∼5-15%) across the entire polyQ tract. Further, visualization of the secondary structure variation over three independent trajectories (∼15 μs each) reveals several instances of stable α-helix formation (>1 μs) across the polyQ tract which was independent of N17 α-helicity (**Fig S3B**) and not observed in case of the AMBER03ws ensemble (**Fig. 1D, S2A**). Finally, the ensemble SS-map indicates weaker structural cooperativity between N17 and polyQ regions for α-helix propagation across these regions (**Fig. S3B, C**), in stark contrast to AMBER03ws (**Fig. 1E**). Overall, we conclude that AMBER03ws provides a more accurate description of the N17-Q16 structural ensemble than AMBER99SB-disp and hence, was subsequently chosen to generate the simulation ensembles of all other constructs presented in this study.

### Emergence of β-sheet conformations upon polyQ length expansion is driven by weak N17- polyQ interactions

*In vitro* experiments indicate that the critical nucleus size for polyglutamine aggregation reduces from a multimer (2-4) to monomer in the case of longer fragments (>Q_25_).^27^ Based on these observations, an outstanding question is how polyQ expansion influences the structural ensemble of the Httex1 monomer and induces conformational changes in the polyQ tract to favor aggregation. Current evidence suggest that the formation of β-hairpin structure in expanded Httex1 strongly favors amyloid aggregation. Firstly, a monoclonal antibody (3B5H10) recognizes a two-stranded β-hairpin conformation in the polyQ tract of mutant Httex1 monomer and its formation *in vivo* effectively predicts neurodegeneration.^62,63^ Secondly, the aggregation rate of a non-pathogenic Httex1 fragment can be enhanced to levels comparable to that of a pathogenic fragment through the introduction of short β-hairpin-forming motifs into polyglutamine sequences.^64^ Despite the importance of β-hairpin conformations in Httex1 aggregation and toxicity, NMR and FRET experiments indicate an increase in α-helicity and rigidification of the polyQ tract with length expansion.^49,56,65,66^ Further, the elevated helicity in the pathogenic Httex1- Q46 also correlated with an enhanced rate of aggregation *in vitro* and *in vivo.*^49^

To address the underlying interplay between α-helix and β-structure formation in the context of polyQ expansion, we performed multi-microsecond simulations (**Table S1**) of N17- polyQ fragments with increasing polyQ length: Q_24/32/46_ and compared their structural ensembles. Consistent with NMR experiments, the per-residue α-helix fractions increased in a polyQ length-dependent manner (Q_16_ to Q_46_) (**Fig. S4A**). The elevated α-helix fraction was predominantly associated within residues beyond the central polyQ tract, implying long-range structural cooperativity across the N17 and expanded polyQ. In the case of N17-Q_46_, however, multi-microsecond simulations (aggregate duration ∼ 105 μs) yielded an ensemble with lower (15-20%) α-helicity per-residue in the polyQ tract compared to NMR estimates (**Fig. S4A**) despite the choice of diverse initial (coil and helical) conformations and convergence of R_g_ distributions across the two sets of trajectories (**Fig. S4B**). With parallel tempering (PT-WTE) simulations (750 ns per replica), we obtained a well-converged ensemble (**Fig. S4C, D**) improved agreement with NMR for α-helicity within the polyQ tract (**Fig. 2A**). These observations highlight the difficulty associated with adequate sampling of structural transitions for IDPs with α-helical propensity using multi-microsecond simulations, due to long helix nucleation times.^67^ The computed SS map of the N17- Q46 ensemble from the PT-WTE temperature replica (293.15 K) clearly indicates the elevated populations of longer helices (25-30 residues) which formed via propagation from the central region as observed for N17-Q_16_ (**Fig. 1E**).

**Figure 2.**
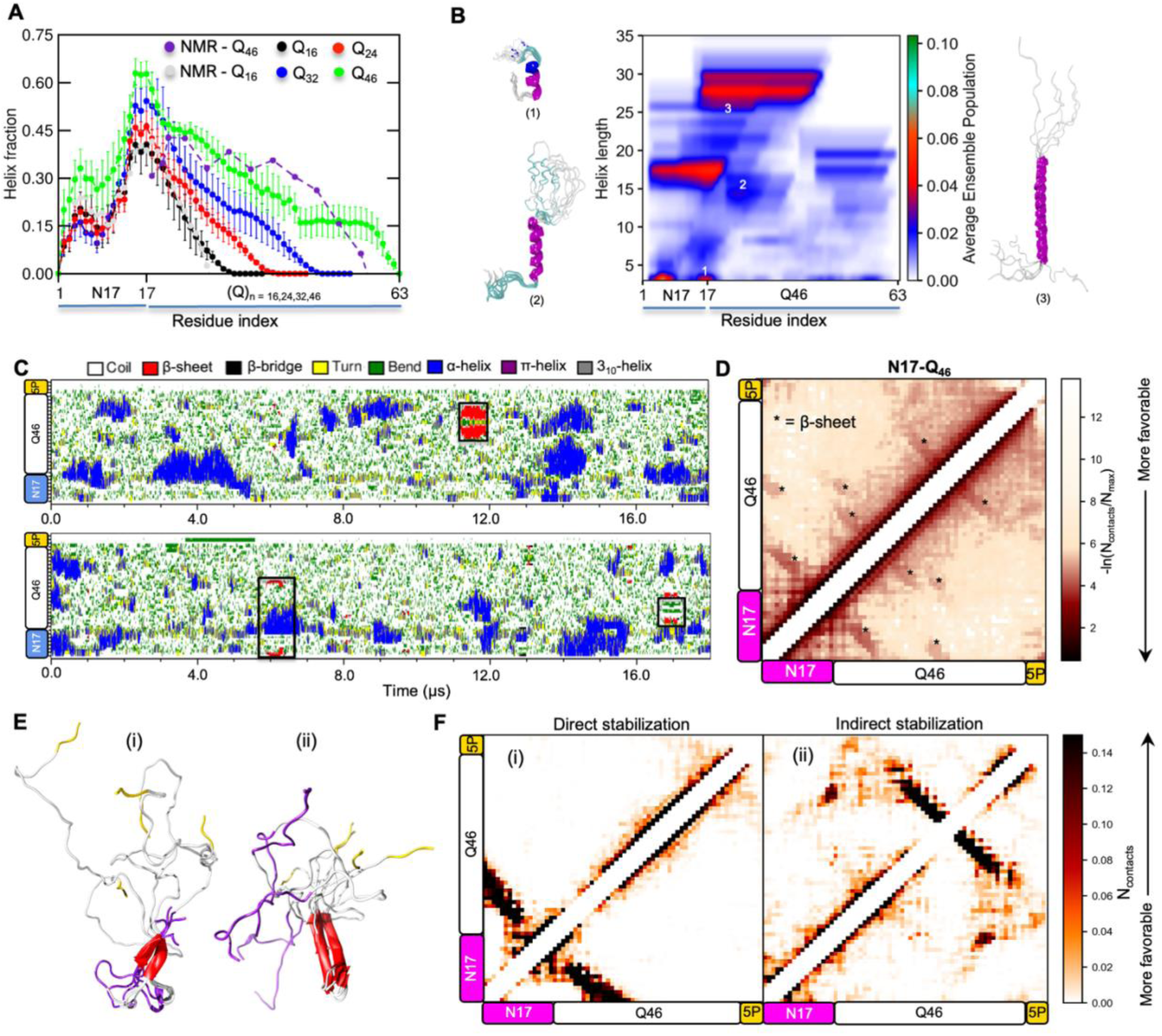
Conformational ensemble of N17-Q_46_. **A.** Comparison of per-residue α-helix fractions for N17-polyQ_n_ constructs with increasing polyQ length (Q_16-46_) against NMR-derived helical propensities (SSP scores). Error bars denote std. error of mean calculated over three independent trajectories for N17-Q_16-32_ constructs and over four intervals (150 ns each) of the PT-WTE replica trajectory (293 K) for N17-Q46. **B.** SS-map of N17-Q46 wild-type (left) from the PT-WTE aggregate trajectory (600 ns) indicating the probability of α-helices formed of varying lengths across N17 and polyQ regions. Representative helical conformations corresponding to numbered regions of the SS-map are shown in cartoon representation. **C.** Secondary structure variation as a function of time for two representative N17-Q_46_ trajectories showing the formation of two-stranded β-sheet structures (black boxes). **D.** Two-dimensional intramolecular contact maps calculated over six N17-Q_46_ independent trajectories indicating the low population of β-conformations (marked as *) relative to α-helices. **E**. Representative β-sheet structures (5 in total) from the trajectory periods highlighted in C (black boxes) involving either direct (i) and indirect (ii) contacts between the N17 and Q46 domains. The coloring scheme for the structures is as follows: N17 - purple, Q46 - white and 5P - Gold, β-strand - red. **F.** Contact maps calculated for the two β-sheet structural ensembles shown in **E.**

A closer inspection of the independent trajectories revealed the formation of transient two- stranded β-sheets (See Methods) for all three expanded N17-polyQ constructs (**Fig. 2C, S5, S6**) which was not observed in N17-Q_16_ and visualized in the two-dimensional intramolecular contact maps computed for each of the ensembles (**Fig. 2D, S7**). For N17-Q_24,_ a two-stranded β-sheet structure formed in two trajectories (total population∼0.5%) (**Fig. S5A**) while only a β-bridge structure involving N17 and polyQ was also observed in another trajectory. For N17-Q_32_, two out of three trajectories showed the formation of two-stranded β-sheet structures (total population ∼ 1.8%) **(Fig S5B)**. Among the six N17-Q46 trajectories, β-sheet conformations involving the polyQ tract were observed in five trajectories (six conformations) with an aggregate population of 1.8% (**Fig. 2C, S6A-B**). Interestingly, half of the β-conformations of N17-Q_46_ were directly stabilized by hydrogen-bonding with N17, the other half involved hydrogen bonding within the polyQ region (**Fig. 2E-F, S5C**). Similar to the unbiased N17-Q_46_ trajectories, a low population of β-sheet conformations (1.1%) were also observed in the PT-WTE 293 K replica trajectory (**Fig. S8**), further confirming their unstable nature compared to α-helical and coil conformations.

The direct participation of N17 in the formation of β-bridges/sheets (**Fig. 2F, left**) raises a possibility that emergence of β-sheet conformations in N17-Q_46_ may be enhanced through transient interdomain interactions. In support of this idea, the contact map of a β-sheet conformation stabilized within the polyQ region also showed weak N17-polyQ interdomain interactions (**Fig. 2F, right)**. As stated earlier, the conformation-specific Httex1 antibody (3B5H10) selectively binds to a two-stranded β-hairpin conformation formed within polyQ tract of pathogenic Httex1.^63^ Notably, the antibody bound to the polyQ tract *in vivo* only when either of the Httex1 flanking domains (N17/PRD) were present.^23^ These observations suggest that polyQ flanking regions such as N17 can modulate the polyQ conformational ensemble and promote the emergence of pathogenic conformations such as intramolecular β-hairpins which favor amyloid aggregation.^64^ To directly test the idea that N17 promotes β-conformations in the polyQ tract through interdomain interactions, we performed multi-microsecond (triplicate) simulations of a simple Q_46_ polypeptide for comparable duration to N17-Q_46_ . The analysis of secondary structure variation over the time course of the Q_46_ trajectories (**Fig. S9**) indicated an absence of β- sheet/bridge conformations which were stable on timescales comparable to those seen for N17- Q_46_, thereby confirming the ability of interdomain N17-polyQ interactions to promote transient chain compaction and form potentially “toxic” β-conformations.

### The interplay between inter-domain interactions and increased polyQ tract α-helicity modulates the oligomerization landscape of Httex1

NMR studies indicate that the tetramerization of Httex1 via the helical N17 region is essential for efficient nucleation and fibrillation.^39^ Based on PRE measurements and simulated annealing,^39^ a structural model of α-helical N17 tetramer (<1% population) was determined for a N17-Q7 construct (high solubility) and found to comprise of two anti-parallel N17 dimer units stabilized by hydrophobic interactions (**Fig. 3A**). The formation of the tetrameric intermediate proceeds through weak dimerization (K_d_ ∼ 0.1-10 mM) as detected by NMR and analytical ultracentrifugation.^39^ The dimeric ensemble is proposed to comprise of “productive” (i.e., anti-parallel) dimers which can effectively assemble into nucleation-competent tetramers and “non-productive” dimers which block tetramerization. Interestingly, the conversion of prenucleation tetramers into amyloid fibrils remains unfavorable for polyQ length < 8 residues.^68^ Therefore, a mere increase in the local concentration of polyQ tracts through oligomerization is insufficient to explain the tetramer- dependent aggregation pathway. Two additional phenomena were invoked based on *in vitro* experiments to explain the underlying mechanism^37^: (i) structural transformation, i.e., an increase in α-helical structure and (ii) domain “cross-talk”, i.e., the emergence of weak inter-domain interactions. However, the precise effects of these two phenomena on the oligomerization landscape of Htttex1 remains challenging to address using experimental techniques such as NMR due to the low population of oligomeric species and poor solubility of Httex1 mutants.

**Figure 3.**
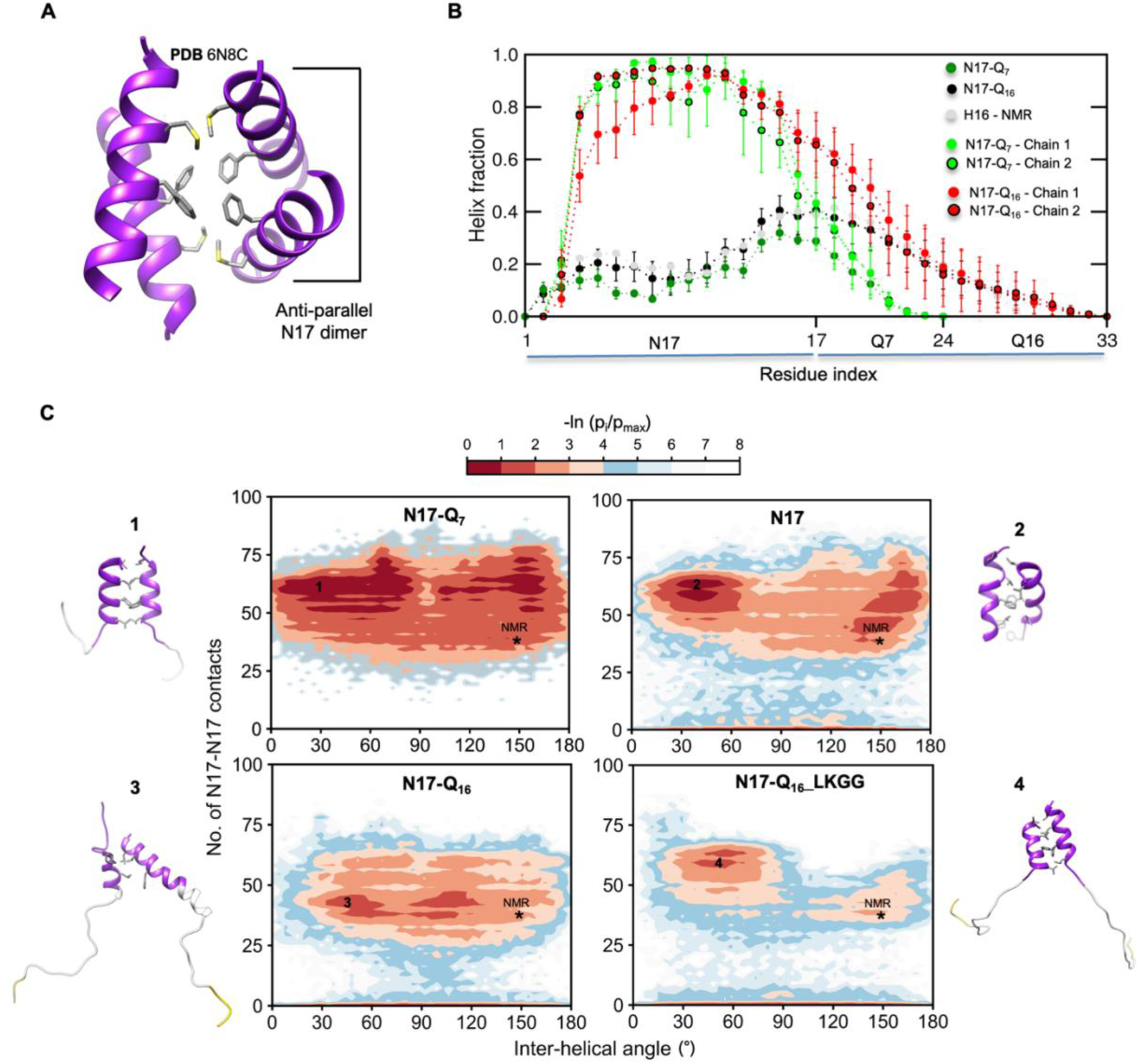
Competition between structural transformation and interdomain interactions shape the dimerization landscape of N17-polyQ. **A.** NMR structure of Httex1-Q7 (H7) tetramer showing interactions (sidechains shown as sticks) in the hydrophobic core. The anti-parallel dimer was extracted from the structure for MD simulations. **B**. Comparison of per-residue α-helix fractions for H7/16 chains from dimer simulations against monomer simulations and NMR. Error bars denote std. error of mean calculated over six independent trajectories. The comparison indicates significantly elevated helicity of N17 region in H7/16 dimers compared to their respective monomers. **C.** Two-dimensional PMF (potential of mean force) plots as a function of two order parameters - (i) total number of intermolecular N17 contacts and (ii) the inter-helical angle (°) between N17 domains, which characterize the association free energy landscape for four N17 dimer variants. The plots shows the combined data derived from 6 independent dimer trajectories (∼2.3 μs) for each variants. The position of NMR - N17 dimer (initial structure) is marked on the PMF plots as (*). Representative dimer MD structures from the basins marked (1, 2) in PMF plots are shown on the left with interfacial hydrophobic residues shown as grey sticks.

In order to tease apart the effects of structural transformation and inter-domain interactions on the oligomerization landscape of Httex1, we first simulated two “productive” dimer variants extracted from the tetramer model as model systems: N17-polyQ_7_ and N17-polyQ_16_. N17-polyQ_7/16_ dimer ensembles showed an oligomerization-dependent stabilization of α-helical structure in N17 **(Fig. 3B, S10A-C)** which is consistent with the CD and NMR experiments.^39,68^ Notably, the expansion of the polyQ tract from 7 to 16 residues also promoted α-helix propagation into the first few glutamine residues, as indicated by their elevated α-helical populations **(Fig. 3B)**. The N17- Q_7_ dimer maintained its stability over microsecond timescale; none of the trajectories showed dissociation of the dimer (**Fig. 3C, S11A** - top row, left) on the simulated timescale (∼2.3 μs per trajectory). To characterize the free energy landscape of the dimer variants, two-dimensional PMF plots were computed based on two order parameters: (i) The total number of contacts between N17 domains of the monomers which distinguishes between stable and weakly-associated states, and (ii) the inter-helical (N17) angle to characterize the diversity of monomer orientation (**Fig**. **3C**). Interestingly, the bound dimer ensemble showed considerable heterogeneity, i.e., in addition to the antiparallel “native” orientation, additional energy basins (favorable regions) were observed for parallel orientations. We speculate that these alternate orientations may be representative of the “unproductive” dimers which are incompatible with higher-order tetramerization.

PolyQ expansion from 7 to 16 appeared to reduce the “native” dimer stability as evident by the loss of numerous native-like and parallel dimer energy basins (**Fig. 3C** - bottom row, left). Three out of six trajectories showed complete dissociation of the dimer followed by multiple weak reassociation events (**Fig S11C**). The secondary dimers showed a reduced number of contacts (<25) between N17 domains and α-helix stability (**Fig. S10B, C**) compared to the bound ensemble. To assess whether the domain “cross-talk” model^37^ could explain the reduced stability of N17-polyQ_7/16_, we computed contact maps of both N17-polyQ_7_/Q_16_ dimer ensembles. In support of the domain “cross-talk” model, contact map analysis indicated an extensive network of inter-domain N17-polyQ interactions for N17-polyQ_16_ (**Fig. S11E, F**).

To determine the effect of increased polyQ helicity on the stability of the bound dimer ensemble, we simulated two dimer variants: N17 and N17-Q_16_ dimer variant with ^14^LKGG^17^ substitution.^55^ The N17 dimer was found to be highly unstable compared to N17-Q_7_; five out of the six trajectories resulted in complete dissociation (**Fig. 3C, 311B** - top row/right) due to the reduced α-helicity of N17 in the absence of a downstream polyQ tract. Similarly, among the ^14^LKGG^17^ trajectories, five out of six trajectories showed complete dissociation followed by weak reassociation (**Fig S11D)**. Accordingly, the resulting dimerization landscape shows a lower population of stable N17-mediated dimers (**Fig. 3C** bottom row, right) and a concomitant increase in the population of weakly-associated dimers. These results support the interpretation that structural transformation (i.e., increase in α-helicity) of the monomer promotes N17 dimer stability and provides a plausible explanation for the reduced aggregation propensity of Httex1-Q_46_ ^14^LKGG^17^ variant as observed in experiments.

The above findings enable us to conclude that an increase in polyQ length can promote transient intra- and intermolecular N17-polyQ interactions which counteracts the stabilizing effect of ɑ-helical propagation into the polyQ tract, thereby promoting a heterogenous dimerization landscape. As a logical extension of this idea, we speculate that for efficient nucleation to occur within the tetrameric intermediate,^39^ polyQ-mediated interdomain interactions (which are enhanced upon length expansion) must ultimately outcompete the stabilizing effect of increased polyQ α-helicity on N17-mediated oligomerization.

### Proline-rich domain (PRD) modulates Httex1 structure and intermolecular interactions within liquid-like assemblies

While the aggregation pathway of Httex1 and a C-terminal construct lacking an intact PRD (P_10_ only) remains identical under *in vitro* conditions^69^, PRD exerts a negative influence on the aggregation kinetics of Httex1 by increasing the relative stability of soluble oligomers compared to amyloid aggregates.^23^ Further, it was recently observed that the presence of PRD strongly promoted the liquid-liquid phase separation^53^ of Httex1-Q_25-97_ under both *in vitro* and *in vivo* conditions when the latter is attached to a fluorescent protein tag^70^ . Interestingly, PRD was also found to suppress the emergence of amyloid aggregates from liquid-like Httex1 condensates (i.e., a “liquid-to-solid” transition).^53^ However, a complete molecular picture of the inter-domain interactions of PRD which contribute towards the stability of liquid-like Htttex1 assemblies is currently unavailable.

To assess the effect of PRD on the structural ensemble of Httex1, we first performed PT-WTE simulations of the Httex1-Q_16_ monomer (**Fig**. **S12**). The ɑ-helical fractions per-residue (**Fig. S12A**) and mechanism of ɑ-helical nucleation in Httex1-Q_16_ are largely similar to N17-Q_16_ (**Fig**. **S12B-C**). These observations collectively indicate that PRD does not impact the structural ensemble of Httex1-Q_16_ which is broadly consistent with ^13^C isotope-labeling NMR experiments wherein the disruption of sequence connectivity between the PRD and polyQ tract (by introducing five glycine residues) only resulted in a modest increase in helicity of the latter^55^. Further, the 2D- intramolecular contact map of Httex1 reveals that intradomain interactions of PRD which involve the polyproline P_10_ and P_11_ tracts are considerably more pronounced compared to interdomain interactions with N_17_ and Q_16_ regions.

To uncover the network of inter-domain interactions implicated in the formation of Httex1 liquid-like assemblies, we performed all-atom simulations of pre-formed N17-Q_16_ and Httex1-Q_16_ condensates (**Fig. 4**). The all-atom condensates were constructed using the phase-coexistence simulation method at coarse-grained resolution and a subsequent backmapping strategy as described previously^71^ (also see Materials and Methods). While the pre-formed Httex1-Q_16_ condensate remained stable over the course of the trajectory, N17-Q_16_ failed to do so (**Fig**. **4A**). Mirroring the interactions observed at the single chain level, analysis of the 2D-intramolecular contact map of Httex1-Q_16_ condensate reveals that homotypic intermolecular interactions between PRD polyproline (P_10_/P_11_) tracts constitute the most significant type of intermolecular interaction (**Fig. 4B**) which clearly highlights the requirement of PRD for condensate formation as seen in experiments. Similar to the case of N17-Q_7-16_ dimers, the ɑ-helicity of the N17 N-terminal region (aa:2-11) is considerably elevated in condensate compared to the isolated monomer due to enhanced intermolecular interactions within the crowed environment of the condensate **(Fig. S13A)**. Further, we also observed the formation of transient intramolecular β-sheets for a population of chains **(Fig. S13B)**. N17 ɑ-helicity of Httex1-Q_16_ is however reduced compared to a uniform N17-Q_16_ condensate system (i.e, without a protein-solvent interface as in the case of Httex1-Q_16_) **(Fig. 4C)**. This implies that N17-PRD interactions **(Fig. 4B)** can suppress the ɑ-helicity of N17 and potentially limit the aberrant interactions which promote liquid-to-solid phase transitions within Httex1 condensates.

**Figure 4.**
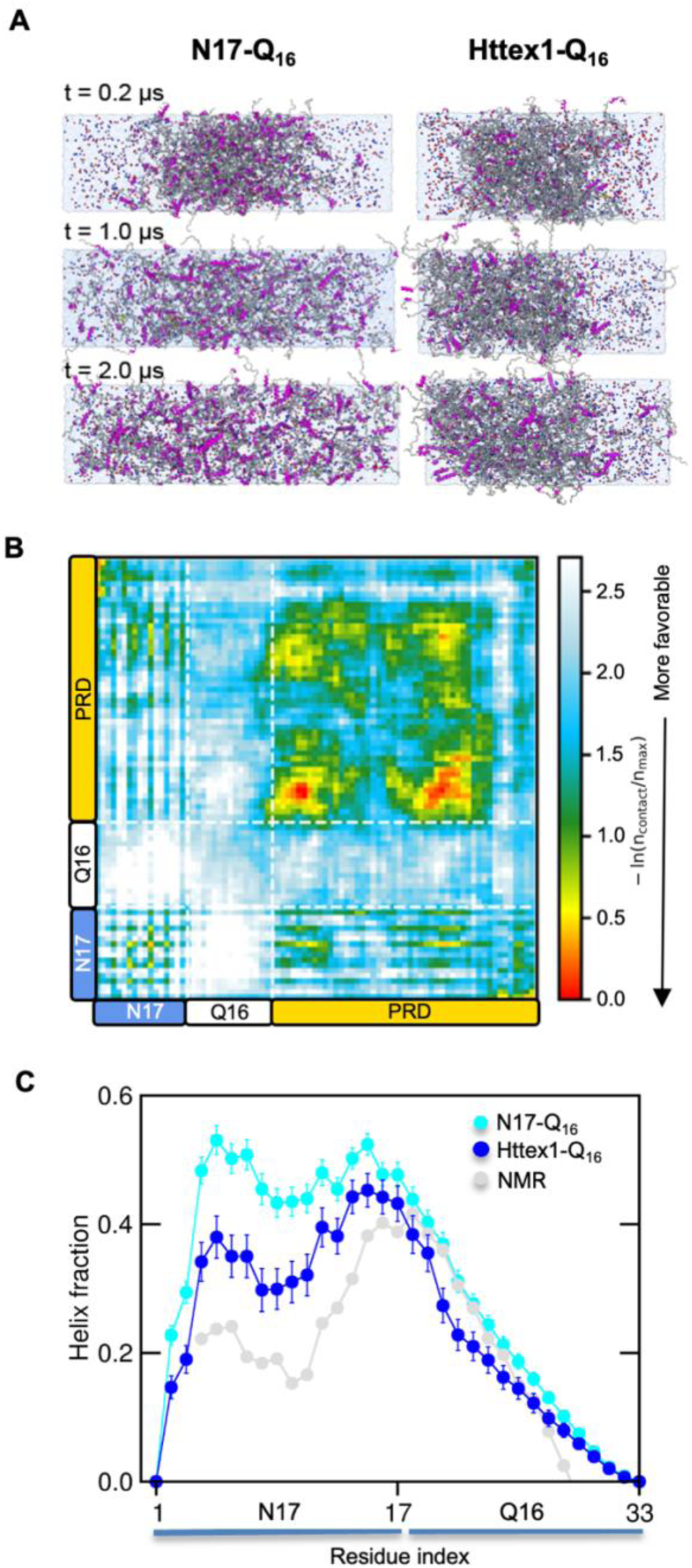
PRD-mediated interactions stabilize the Httex1-Q_16_ condensate and suppress N17 ɑ- helicity. **A.** All-atom MD simulation snapshots of N17-Q_16_ (left) and Httex1-Q_16_ (right) condensate systems over 2 μs of their respective trajectories. Protein chains are show as ribbons and ions (Na^+^/Cl^-^) are shown as spheres. The snapshots indicate destabilization of the N17-Q_16_ dense phase (0.2 - 1.0 μs) and its complete dissolution (2.0 μs), resulting in a homogeneous system. **B.** Pairwise 2D- intermolecular contact map averaged over all chain pairs reveals significant contributions of the PRD towards stabilization of the Httex1-Q_16_. **C.** Mean fractional ɑ-helicity per-residue (computed over all dense phase chains) shows reduced ɑ-helical formation within N17 for the Httex1-Q_16_ condensate compared to a homogenous N17-Q_16_ condensate system (i.e., lacking protein-solvent interface). Error bars denote std. error of mean over all chains.

**Figure 5.**
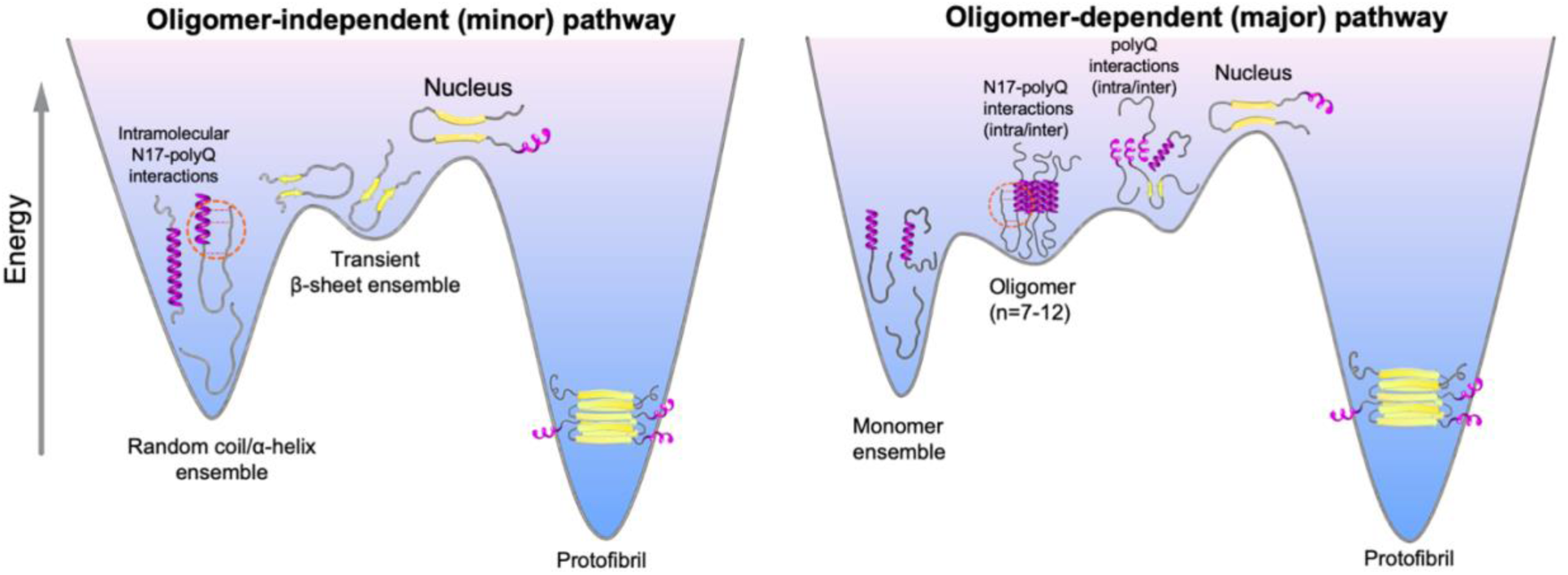
polyQ expansion promotes “domain crosstalk” which modulates the structural ensemble and oligomerization landscape of Httex1 to favor fibrillar aggregation. Schematic illustrating the effect of polyQ length expansion on the conformation and oligomerization of Httex1. polyQ expansion promotes inter-domain N17-polyQ interactions. These interactions can (i) induce the emergence of transient β-sheet conformers in monomers which may fold to form the critical nucleus and initiate protofibril formation (oligomer-independent pathway), and (ii) counteract stabilization of N17-mediated oligomers to favor polyQ-polyQ interactions which lead to nucleation and fibrillation. The nucleus size is shown as a monomer (n=1) and is representative of Httex1-Q_>23_.

## Discussion

Emerging evidence from biophysical investigations suggest that there is a complex interplay between the N-terminal and C-terminal flanking sequence composition and structure, and polyQ length in modulating the overall aggregation propensity of polyQ proteins. Among the polyQ proteins implicated in neurodegenerative disorders, three of the proteins: Huntingtin exon 1 (Httex1), Androgen Receptor (AR), and Ataxin-7 (Atx-7) contain hydrophobic N-terminal flanking sequences which induce and progressively increase the α-helical propensity in the polyQ tract with length expansion.^55,72,73^ However, the increased α-helical propensity in the polyQ region appears to have dissimilar effects on the aggregation of these polyQ-containing sequences. While both AR and Atx-7 showed suppressed polyQ aggregation for polyQ_25-33_,^72,73^ aggregation of longer polyQ sequences (Q_46-72_) is favored in the case of Httex1.^56^ It is therefore important to decipher the role of polyQ flanking sequences in order to obtain mechanistic insights into the amyloid aggregation of polyQ proteins and develop novel therapeutics to treat their associated- neurodegenerative disorders.

The rapid aggregation kinetics of pathogenic Httex1 fragments limit the ability to achieve a complete atomic-level characterization of the role of flanking domains on structural and oligomerization landscape through experimental approaches alone. Further, conflicting results from previous MD simulation studies^74–76^ due to force field and water model inaccuracies has led to a lack of clarity regarding the precise molecular mechanisms of Httex1 oligomerization and pathogenic aggregation. Our study provides a comprehensive characterization of the structural ensembles of normal and pathogenic N17-polyQ fragments using the AMBER03ws force field which was carefully tuned to achieve balanced secondary structure propensities and protein-water interactions.^47,77^ Importantly, our simulation ensembles correctly captured the length-dependent increase in helical from Q_16_ to Q_46_, achieving good quantitative agreement with NMR in the case of N17-Q_16/46_ fragments. Therefore, a distinguishing feature of this computational study is the extensive structural validation of N17-polyQ simulation ensembles against recent high- resolution NMR experiments,^55,56^ which considerably enhances the reliability of our predictions. In addition, our simulations also captured the effect of point mutations in the hydrophobic four-residue motif (^14^LKSF^17^) which structurally couples N17 with the polyQ tract^55^. The N17 domain is also subject to numerous post-translational modifications (PTMs) which alter its α-helical structure and impacts the cellular localization, oligomerization, and aggregation of Htt.^78–81^ Our chosen model and simulation methodology demonstrated here opens the way for accurate predictions of the effect of point mutations and PTMs on the structure and conformation of Httex1 and other polyQ proteins which contain N-terminal hydrophobic motifs adjacent to the polyQ tract.

Two opposing models are proposed to explain the coupling between structure and binding properties of polyglutamine with increasing length, namely the (i) “linear lattice” model and, (ii) “conformational emergence” model. The linear lattice model proposes that the enhanced multivalency (number of interaction sites) of polyglutamine upon length expansion coupled with the prevalence of an extended, random coil structural ensemble (abundant in N17-Q_46_/Q_46_ simulations) explains its aberrant cellular interactions and increased binding affinity towards anti-polyQ antibodies.^82–84^ Nevertheless, several studies also point towards the existence of β-hairpin conformations formed in pathogenic polyglutamine^17^ and Httex1^23,62,85,86^ in monomeric and oligomeric species. Interestingly, our results appear to reconcile the two opposing models in the case of pathogenic Httex1, by suggesting that the increased multivalency can also promote intramolecular interdomain (N17-polyQ) interactions which may give rise to “toxic” β- conformations or those which proceed to form the aggregation-competent nuclei more readily compared to disordered monomers (**Fig. 5**). Alternatively, such interactions may also promote polyglutamine “folding” to form the critical β-hairpin nucleus, which can template the formation of amyloids through a kinetically competing N17-independent pathway.^33^ Overall, our simulations directly detect the emergence of a conformational equilibrium between the predominant disordered conformer and potentially “toxic” β-sheet conformers upon polyQ expansion, thereby providing a critical insight into polyQ aggregation and toxicity. It is important to note, however, that β-conformers observed in our N17-Q_32/46_ simulations constitute a lowly populated species (1- 2%), implying that such conformations are highly unfavorable relative to helical and coil conformations. These observations are qualitatively consistent with a highly unfavorable free energy of folding associated with polyglutamine nucleus formation (+12.2 kcal mol^-1^ for Q_47_).^26^

We provide a high-resolution view into the competing effects of domain “cross-talk” and structural “transformation” on Httex1 oligomerization, by directly demonstrating that^35^: (i) interdomain N17-polyQ interactions destabilize oligomerization and, (ii) α-helical propagation into the polyQ tract which stabilizes oligomerization. These observations lead us to conclude that polyQ-mediated interdomain interactions must outcompete the stabilizing effect of increased polyQ α-helicity to favor nucleation and conversion of oligomers into amyloid aggregates. Our simulations of the Httex1 monomer and condensates uncover the significance of intermolecular PRD interactions in driving the formation of liquid-like Httex1 assemblies. Further, PRD-N17 intermolecular interactions which were shown to suppress N17 helicity could also limit the formation of aggregation-competent oligomers and amyloid aggregates within Httex1 condensates. In conclusion, our findings support the idea that altering the balance of homo- (N17)^87^ and heterodomain (N17-polyQ/PRD) interactions^32^ may serve as an effective therapeutic strategy for the inhibition of pathological self-assembly of Httex1.

## Acknowledgments

This work was supported by NIH/NIGMS R35GM153388. Atomistic MD simulations were conducted with the advanced computing resources provided by Texas A&M High Performance Research Computing. We are thankful to Pau Bernado and Amin Sagar for providing us with the ^13^C NMR chemical shift data to validate the atomistic simulations presented in this work and their critical feedback on the manuscript during its preparation. We would also like to gratefully acknowledge Randal Halfmann and Shriram Venkatesan for fruitful discussions, and their critical feedback during the initial writing process and preparation of the final manuscript.

## Materials and Methods

### Initial structure, force field choice and system setup

For N17-polyQ_16/24/32_ monomer, simulations were initiated from three different coil configurations generated based on the Flexible Meccano algorithm^88^ using the ProtSA webserver.^89^ For PT-WTE simulations N17-polyQ_46_, a single coil conformation was chosen among the three used for unbiased simulations. The three partially-helical, initial conformations of N17-Q_46_ were obtained from the first microsecond of a coil trajectory. For dimer simulations, the initial complex was extracted from the lowest energy NMR model of the N17 tetramer (PDB 6N8C). PolyQ sequences (Q_7/16_) were added to the monomer units of the N17 dimer using MODELLER.^90^ The initial configurations were solvated in octahedral boxes with 150 mM NaCl, and additional counter ions added to achieve electroneutrality. The polypeptide and water topologies were described using parameters from the AMBER03ws force field^47^ (https://bitbucket.org/jeetain/all-atom_ff_refinements/src/master/) and TIP4P/2005 water model^91^ respectively. For Na^+^ and Cl^-^ ions, modified LJ parameters^92^ were used to improve ion solubility. Details of box dimensions and duration of simulation trajectories for each N17-polyQ monomer and dimer trajectories are provided in **Table S1**.

### Conventional MD Simulation Protocol

To remove any steric clashes, the solvated polypeptide systems were first relaxed by performing energy minimization using the steepest descent algorithm in GROMACS-2020.4.^93^ Following minimization, the systems were simulated for 100 ps using Nose-Hoover thermostat^94^ with a coupling constant (τ_c_) of 1 ps to achieve temperature equilibration at 293.15 K. Next, a 100 ps simulation was conducted using the Berendsen barostat^95^ with isotropic coupling and a coupling constant (τ_p_) of 5 ps for pressure control, to achieve pressure equilibration at 1 bar. Production simulations were performed in the NVT ensemble was performed using the Langevin Middle Integrator^96^ (friction coefficient = 1 ps^-1^) in OpenMM-7.5.^97^ Short-range nonbonded interactions were computed using a cutoff radius of 0.9 nm, while long-range electrostatics were described with the PME method.^98,99^ The hydrogen masses were increased by 1.5 times to enable a simulation timestep of 4 fs^100^ for the production runs, and hydrogen-containing bonds were constrained using the SHAKE algorithm.^101^

### Parallel Tempering Well-Tempered Ensemble (PT-WTE) MD Simulation Protocol and Assessment of Convergence

Following energy minimization, temperature, and pressure equilibration of the system at 293.15 K and 1 bar, temperature replicas were prepared and equilibrated in the NVT ensemble without exchange for 10 ns at higher temperatures using the Langevin integrator (friction coefficient = 1 ps^-1^). A total of 16 temperature replicas were generated based on a geometric progression that spanned a range from 293 to 500 K. Long-range electrostatic interactions were modeled using the Particle Mesh Ewald method^98,99^ with a real space distance cutoff of 0.9 nm. The hydrogen mass repartitioning scheme^100^ was used to reduce the computational cost along the LINCS algorithm^102^ which was used to constrain all bonds, allowing a time step of 5 fs. Following temperature equilibration of the replicas, production runs were performed in the NVT ensemble with bias potentials (bias factor = 72 for N17-Q_46_ / 96 for Httex1-Q_16_) deposited every 4 ps to increase fluctuations in the system potential energy (collective variable) resulting in a well-temperature ensemble (WTE) with improved frequency of conformational exchange between adjacent replicas,^103,104^ which was attempted every 1 ps. The initial height and width of the Gaussian potential were set to 1.5 and 456.0 kJ mol^-1^. Exchange probabilities between adjacent replicas varied from 17-30%.

For N17-Q_46_, the first 150 ns from the 293 K replica was discarded as equilibration time respectively, during which the initial height of the gaussian potential dropped below 0.25 kJ mol^-1^, and sufficient overlap in the potential energy distributions between adjacent temperature replicas was observed. The initial configurations for each replica underwent several round trips through the entire temperature space, which indicates efficient conformational exchange between replicas. The convergence of the 293 K replica trajectory was assessed by computing the radius of gyration (R_g_) and ɑ-helical fraction over independent blocks of the trajectory. **(Fig. S4C,D)**.

### All-atom condensate simulations

All-atom dense-phase simulations of N17-Q_16_ and Httex1- Q_16_ were prepared using the method previously.^71^ Initial structures for multichain simulations were generated by equilibrating 170 chains of N17-Q_16_ at 200 K and 90 chains of Httex1-Q_16_ at 300 K, using coarse-grained (CG) coexistence simulations with the HPS-SS model.^105,106^ Notably, N17- Q_16_ was unable to form a stable condensate at 300 K under these simulation conditions. Subsequently, the CG dense phase configurations in the slab geometry were used to reconstruct all-atom slab configurations with Modeller.^90,107^ Potential steric clashes were resolved through short molecular dynamics simulations with OpenMM-7.6^97^ employing the AMBER03^108^ force field and OBC implicit solvent model.^109^ The resulting relaxed protein systems were then subjected to equilibration and production simulations as described in the preceding sections. The resulting relaxed protein systems were then subjected to equilibration in Gromacs as described in the preceding sections. The Gromacs files were converted to Amber inputs using the “gromber” function in Parmed^110^ and hydrogen mass repartitioning^111^ to 1.5 amu was performed during conversion to facilitate a 4 fs time step. Production simulations were performed using the AMBER22 simulation package.

### Trajectory Analysis

Secondary structure calculations were performed based on the DSSP library^112^ using the gmx do_dssp program. The calculation of NMR chemical shifts and comparison with experiments was performed using SPARTA+ as described in our previous study^113^. The calculation of secondary structure (SS) maps for assessing the α-helicity cooperativity of N17-polyQ regions was performed using a modified version of the python script provided as part of the original study to include the DSSP definition for identification of α-helices. For the detection of bifurcated hydrogen bonds (B-HBs), gmx hbond in combination with in-house python scripts were used to compute the fraction of trajectory frames in which glutamine sidechain to backbone (i → i-4) hydrogen bonds coexisted with backbone-backbone (i-4 → i) hydrogen bonds. B-HB analysis was carried out only on frames in which >30% of the residues adopted an α-helical conformation. For the analysis and visualization of β-sheet conformations, only frames where >4 residues adopted a β-sheet conformations were extracted from unbiased and PT-WTE trajectories. For contact analysis, two residues i and j with sequence separation greater 3 (|i-j|>3) were in contact if any two heavy atoms of the residues were within 0.6 nm of each other. The chosen distance cutoff has been extensively tested in our previous studies^71,113,114^ and is large enough to include the diverse contact modes such as hydrogen bonds (<0.35 nm), van der Waal interactions and salt-bridges (<0.6 nm). For the analysis of N17-polyQ dimerization, the minimum distance, and orders parameters (number of N17-N17 contacts and inter-helical angle) for 2D- PMFs **(Fig. 3C)** were computed using gmx mindist and gmx gangle.

