## Supporting Information for "Transient interdomain interactions modulate the monomeric structural ensemble and self-assembly of Huntingtin Exon 1"

| N17-polyQ construct | Box length (nm) | No. of replicates | Aggregate duration (μs) |
|---|---|---|---|
| N17-$Q_{16}$ - $^{14}$LKSF$^{17}$ - 5P | 8.0 | 3 | 61.9 |
| Ac - $Q_{16}$ - 5P | 8.0 | 3 | 62.1 |
| Ac - $Q_{16}$ - NMe | 8.0 | 3 | 62.1 |
| N17 - $Q_{16}$ - $^{14}$LKAA$^{17}$ - 5P | 8.0 | 3 | 41.1 |
| N17 - $Q_{16}$ - $^{14}$LLLF$^{17}$ - 5P | 8.0 | 3 | 64.2 |
| N17 - $Q_{16}$ - $^{14}$LKGG$^{17}$ - 5P | 8.0 | 3 | 64.7 |
| N17 - $Q_{24}$ - 5P | 8.5 | 3 | 55.4 |
| N17 - $Q_{32}$ - 5P | 9.0 | 3 | 63.3 |
| N17 - $Q_{46}$ - 5P (unbiased) | 10.0 | 6 | 107.2 |
| N17 - $Q_{46}$ - 5P (PT-WTE) | 10.0 | 16 | 12.0<br>(0.75 per replica) |
| Httex1 - $Q_{16}$ (PT-WTE) | 10.0 | 16 | 5.6<br>(0.35 per replica) |
| Ac - $Q_{46}$ - NMe | 8.0 | 3 | 46.4 |
| N17_dimer | 8.0 | 6 | 13.7 |
| N17 - $Q_7$_dimer | 8.0 | 6 | 13.8 |
| N17 - $Q_{16}$_dimer | 8.0 | 6 | 13.5 |
| N17 - $Q_{16}$_dimer - $^{14}$LKGG$^{17}$ | 8.0 | 6 | 13.9 |

**Table S1. Number of replicas and aggregate duration for N17-polyQ monomers and dimer trajectories generated in this study.**

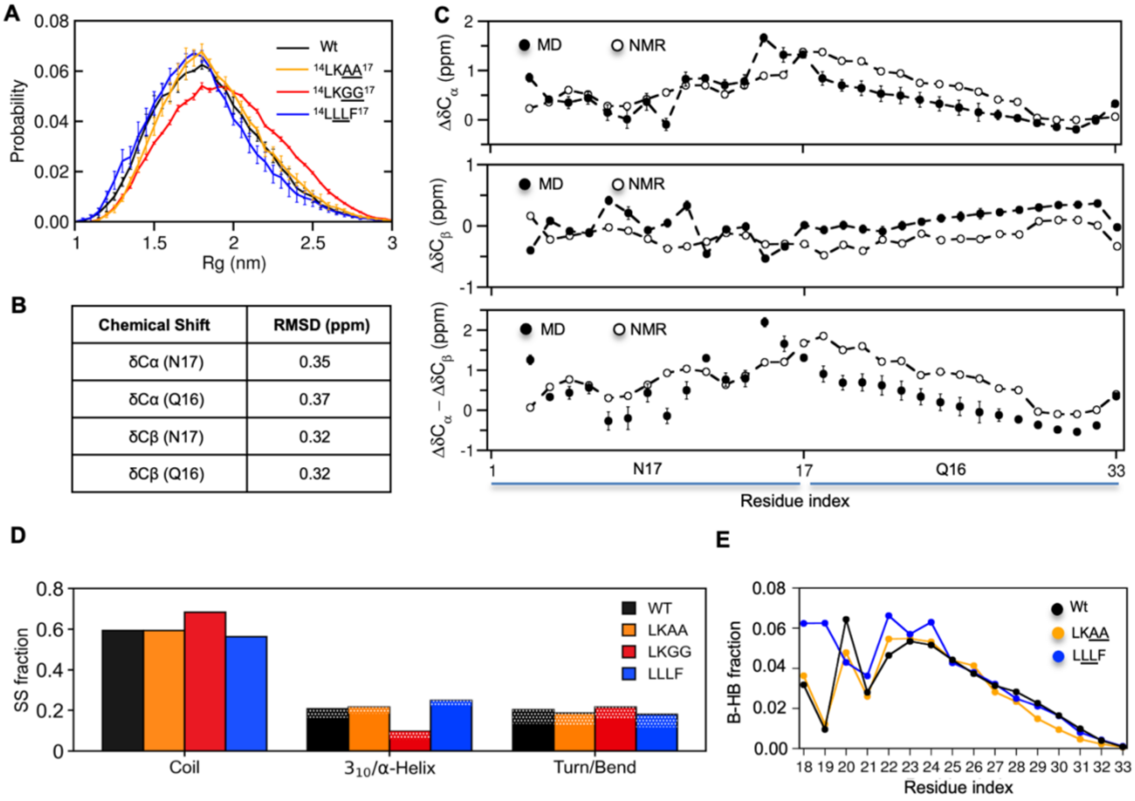


| Chemical Shift | RMSD (ppm) |
|---|---|
| δCα (N17) | 0.35 |
| δCα (Q16) | 0.37 |
| δCβ (N17) | 0.32 |
| δCβ (Q16) | 0.32 |

**Figure S1. Assessment of convergence and validation of N17-polyQ$_{16}$ ensembles against NMR chemical shift data.** Related to Figure 1. **A**. Mean probability distribution for radius of gyration ($R_g$) calculated over three three independent trajectories. Error bars denote std. error of mean. **B**. Chemical shift RMSD (ppm) for predicted values with respect to experiment for $^{13}C$ primary chemical shifts and secondary chemical shifts. Chemical shifts from MD trajectories were predicted using SPARTA+. **C**. Comparison of per-residue secondary chemical shifts with experiment. Error bars denote std. error of mean computed over three independent trajectories. **D**. Mean fraction of total residues involved in the formation of various secondary structure (SS) elements calculated over three independent trajectories for N17-Q$_{16}$ wild type and $^{14}$LKSF$^{17}$ variants. Error bars denote std. error of mean. Dotted regions in the middle and right bar sets correspond to the fraction of residues in $3_{10}$ and turn conformations. **E.** Per-residue fraction of bifurcated hydrogen bonds (B-HB) in the polyglutamine tract computed for N17-Q$_{16}$ wild type and mutants - $^{14}$LKAA$^{17}$/$^{14}$LLLF$^{17}$. For the analysis, only the frames where more than 30% of the protein residues adopted an α-helical conformation were considered.

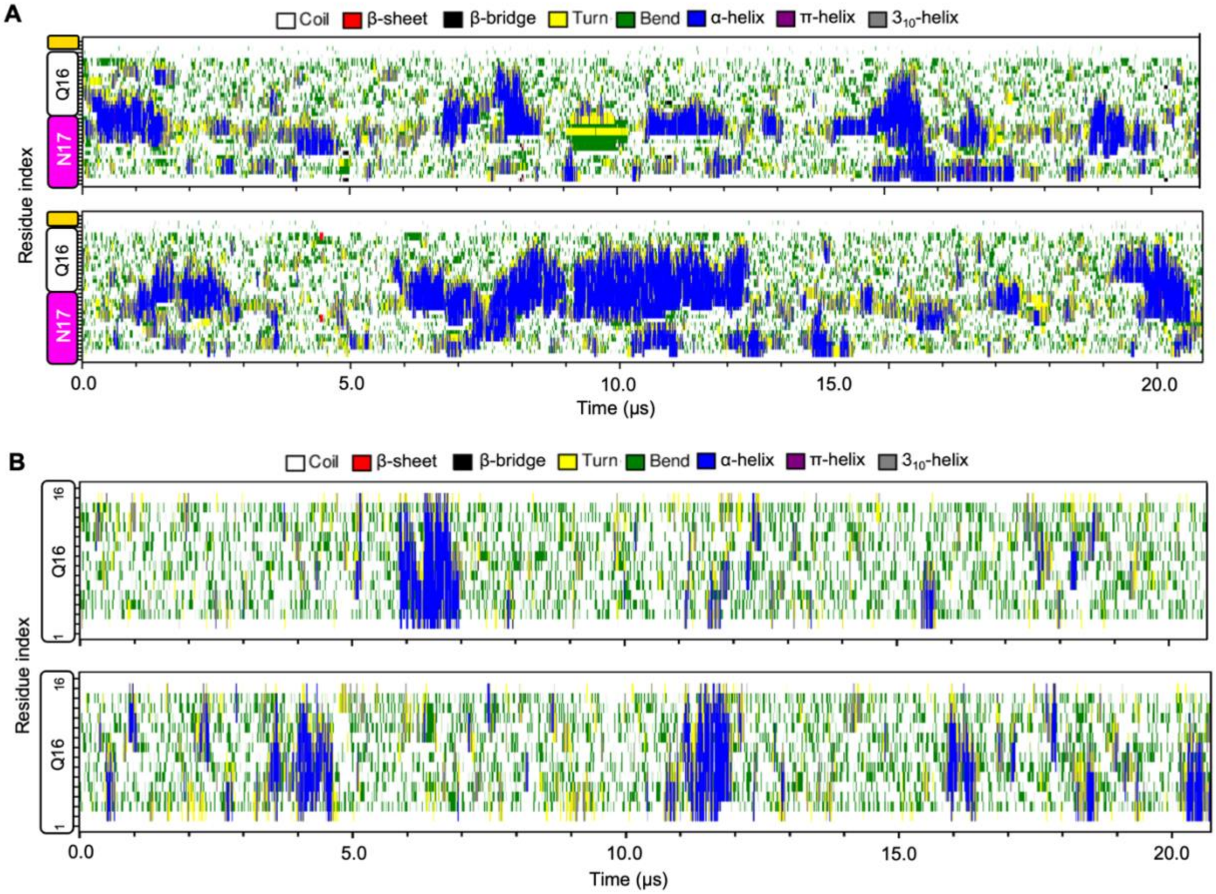


**Figure S2. Dynamics of secondary structure formation in N17-$Q_{16}$ and $Q_{16}$ fragments.** Related to Figure 1.

**A**. Secondary structure variation as a function of time for N17-$Q_{16}$ fragment from two independent trajectories. **B**. Same as in A for Q16.

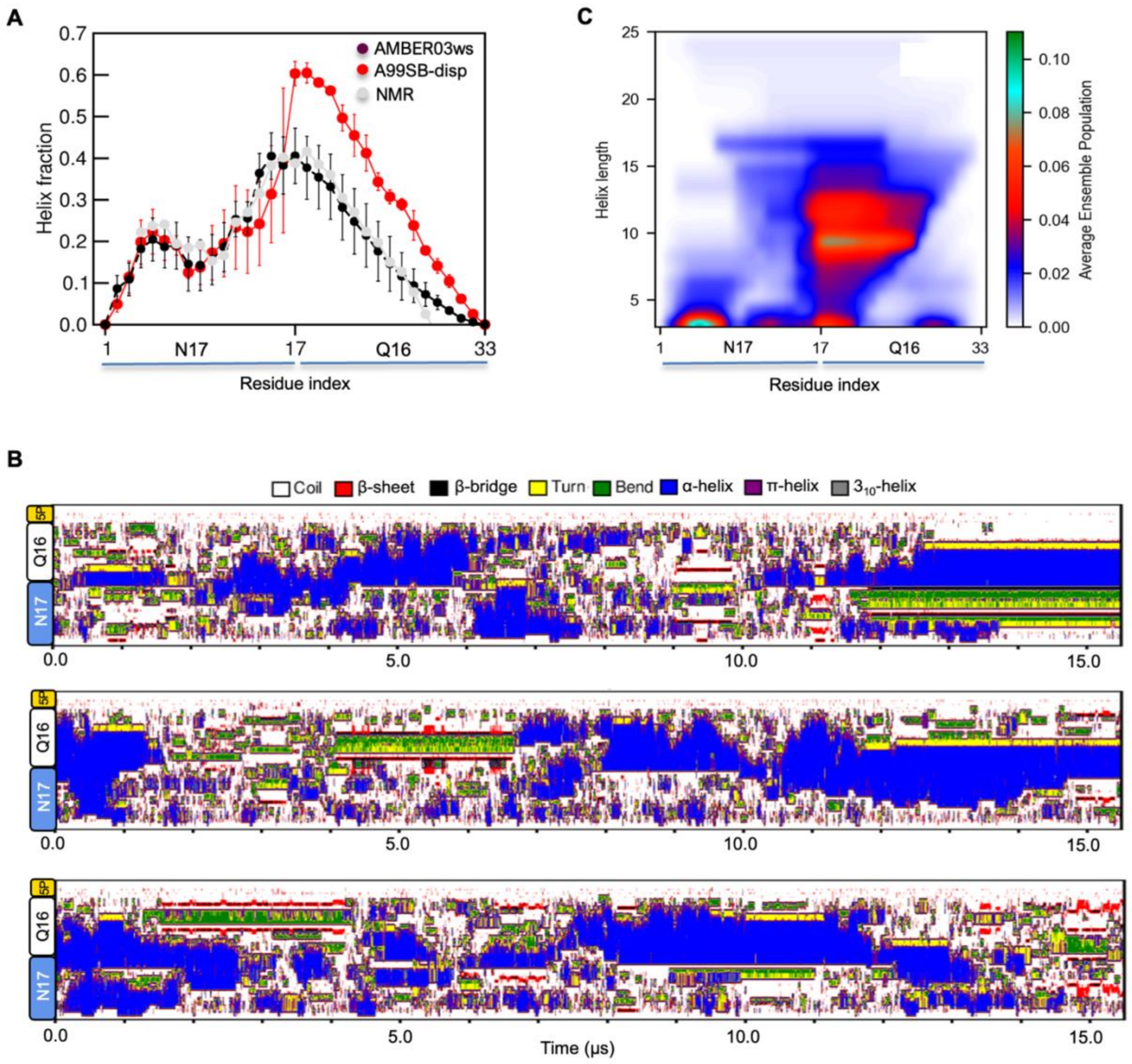


**Figure S3. Structural ensemble of N17-$Q_{16}$ generated using AMBER99SB-disp force field.** Related to Figure 1.

**A**. Comparison of mean α-helical fractions calculated over three independent AMBER99SB-disp/AMBER03ws trajectories initiated from coil conformations and their comparison to NMR SSP scores. Error bars represent SEM over three independent replicate trajectories. **B**. Secondary structure (DSSP) variation as a function of time for each trajectory. **C**. SS-map of N17-$Q_{16}$ wild-type computed from an aggregate trajectory (~45 μs) indicating the probability of various helical lengths across N17 and polyQ regions.

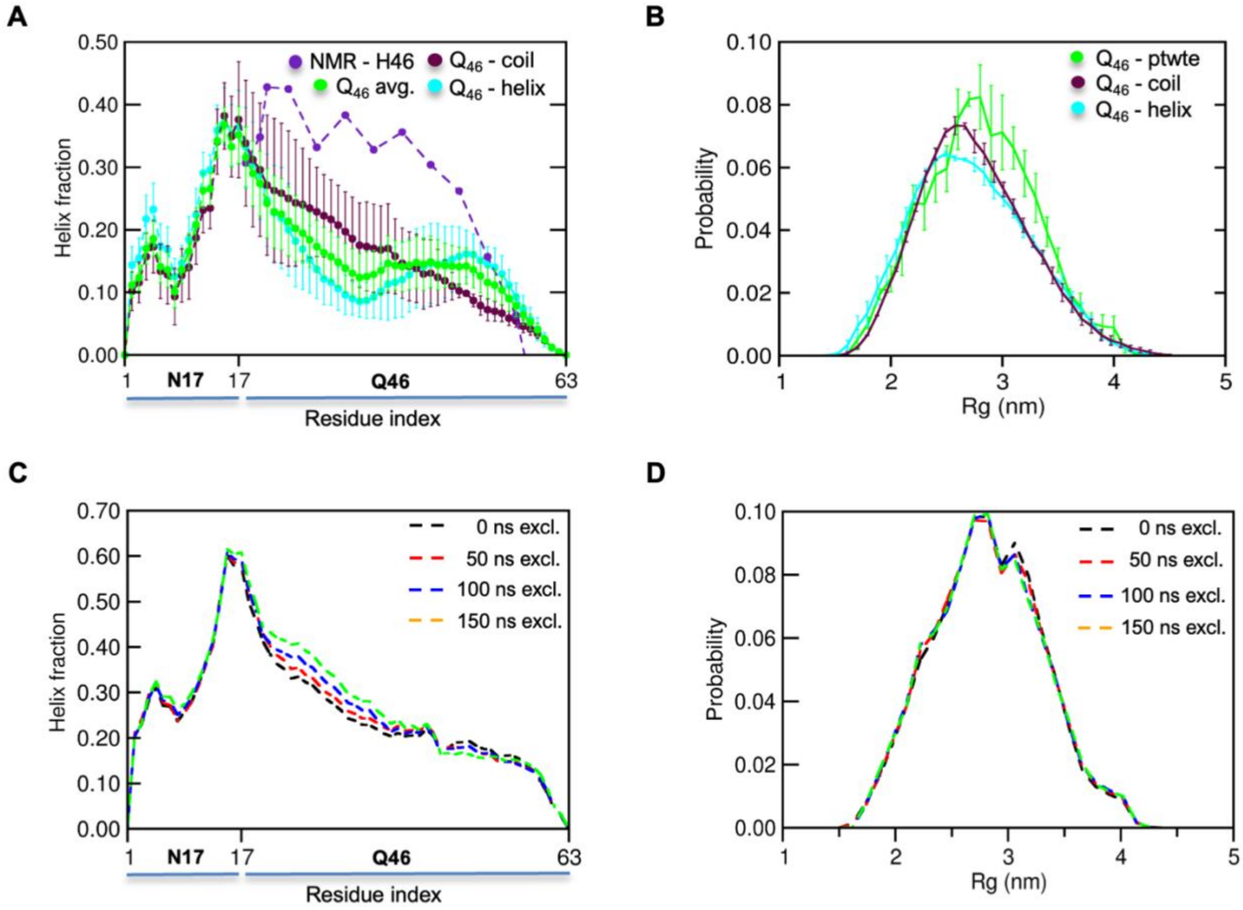


**Figure S4. N17-$Q_{46}$ ensembles generated from unbiased and PT-WTE simulations.** Related to Figure 2.

**A**. Mean fraction of calculated over three independent trajectories initiated from coil and partially-helical conformations. Error bars denote std. error of mean. **B**. Mean probability distributions for radius of gyration ($R_g$) calculated over the unbiased trajectories and the PT-WTE trajectory (500 ns) at 293.15 K. Error bars denote std. error of mean calculated over independent trajectories for coil/helix and 4 blocks (150 ns) of the PT-WTE trajectory. **C.** Per-residue α-helical fractions and **B.** radius of gyration (Rg) probability (right) distributions (dashed lines) of 293 K replica ensembles calculated over the whole 750 ns trajectory (black), first 50 ns (red), first 100 ns (blue) and first 150 ns (green) excluded overlap indicating good convergence of local structure and global dimensions respectively.

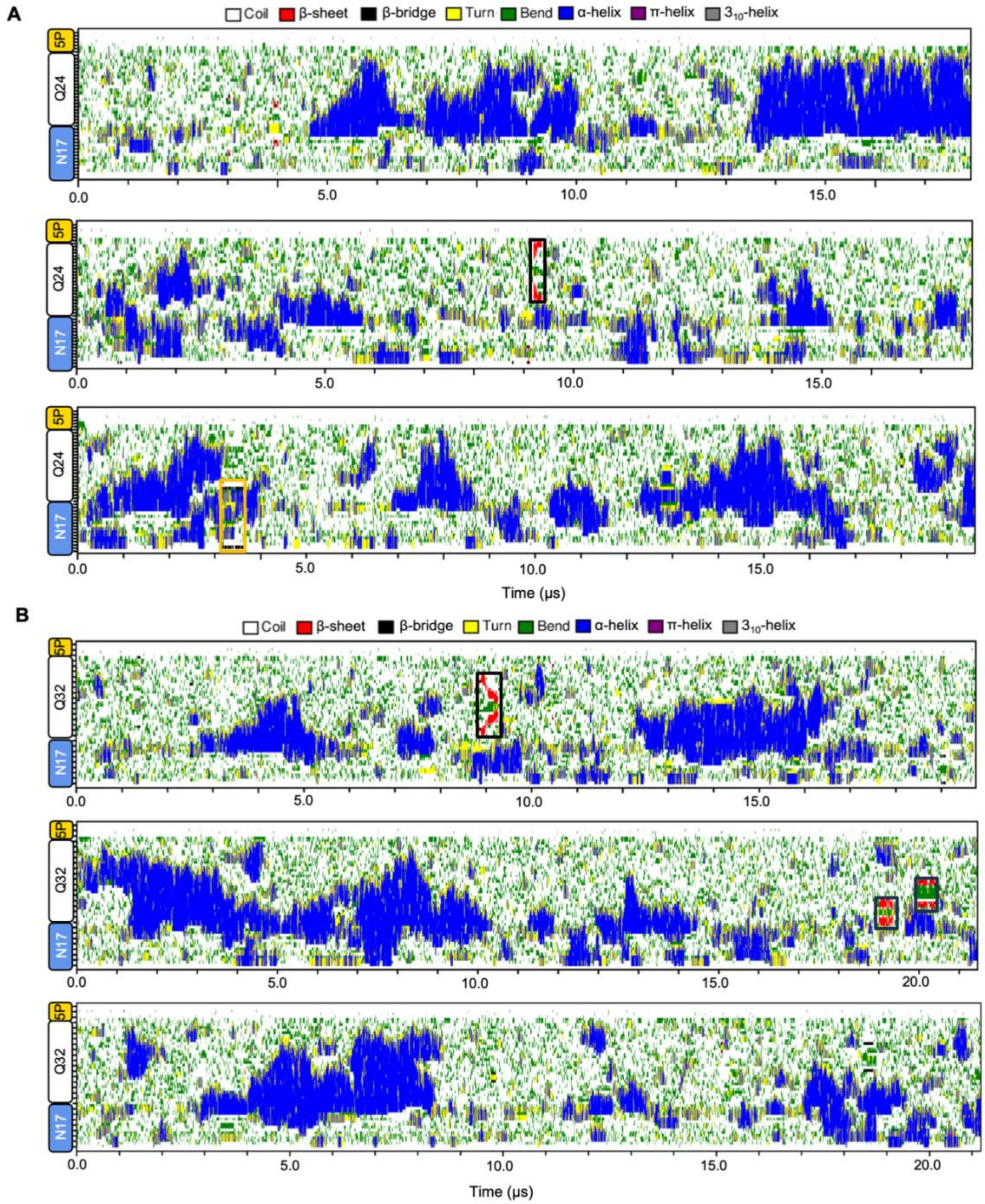


**Fig S5. Secondary structure dynamics in N17-$Q_{24/32}$ trajectories.**
**A.** DSSP secondary structure variation as a function of time for three N17-$Q_{24}$ trajectories initiated from random coil conformation. **B.** Same as in A for N17-$Q_{32}$. In both A and B, β-bridge/sheet structures are highlighted in black boxes.

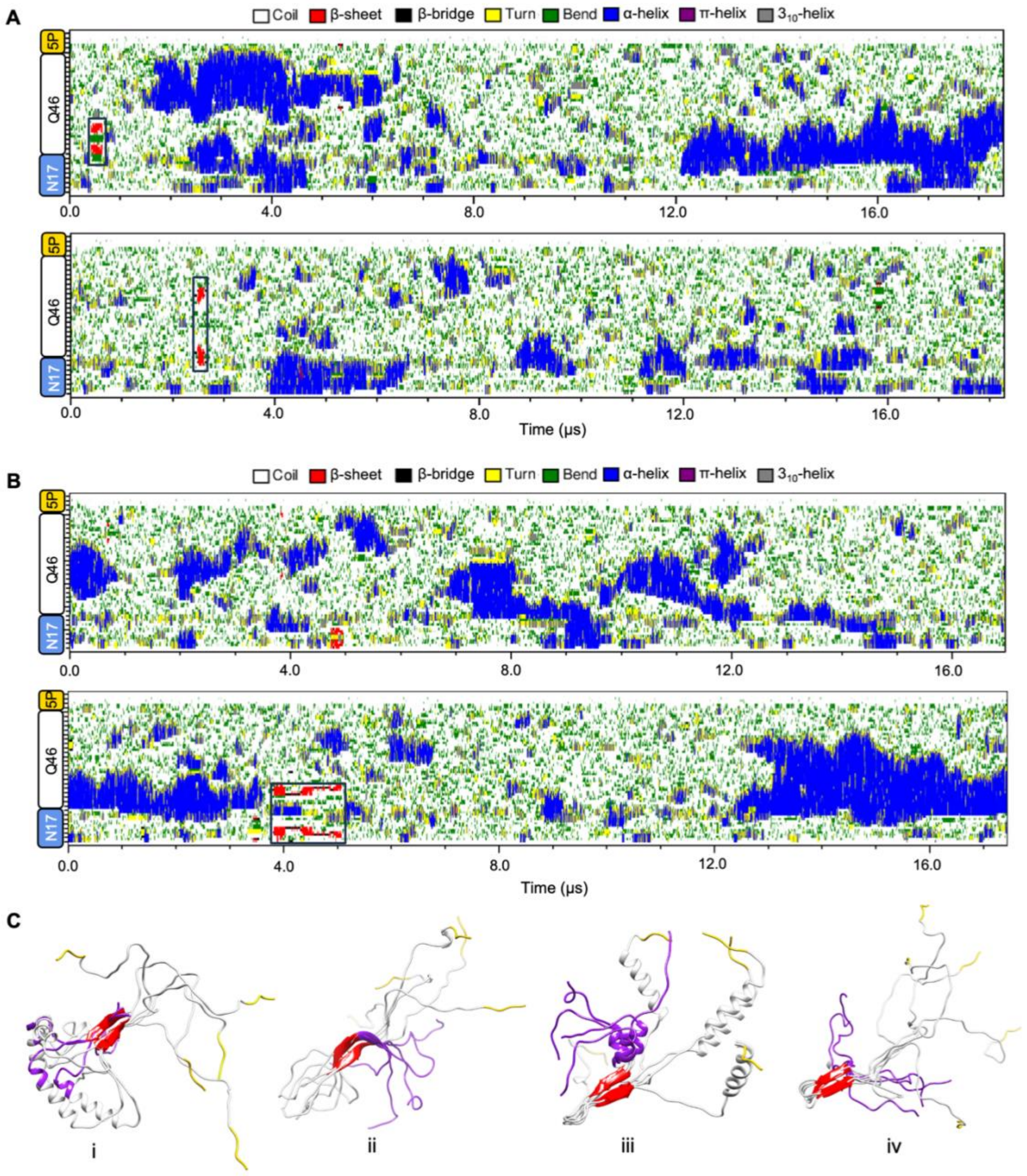


**Fig S6. Secondary structure dynamics in N17-$Q_{46}$ trajectories.** Related to Figure 2.
**A.** Secondary structure variation as a function of time for unbiased H46 trajectories initiated from random coil conformation. β-sheet structures formed are highlighted in black boxes. **B**. Same as in A for two trajectories initiated from partially helical conformations. **C**. β-sheet ensembles (i - iv) from trajectory intervals highlighted in black boxes in panels A/B and Figure 2C. Structures i /ii formed N17-polyQ β-sheets while iii/iv formed intra-polyQ β-sheets. Five representative structures are shown for each ensemble. The coloring scheme for the structures is as follows: N17 - purple, Q46 - white and 5P - Gold, β-strands - red.

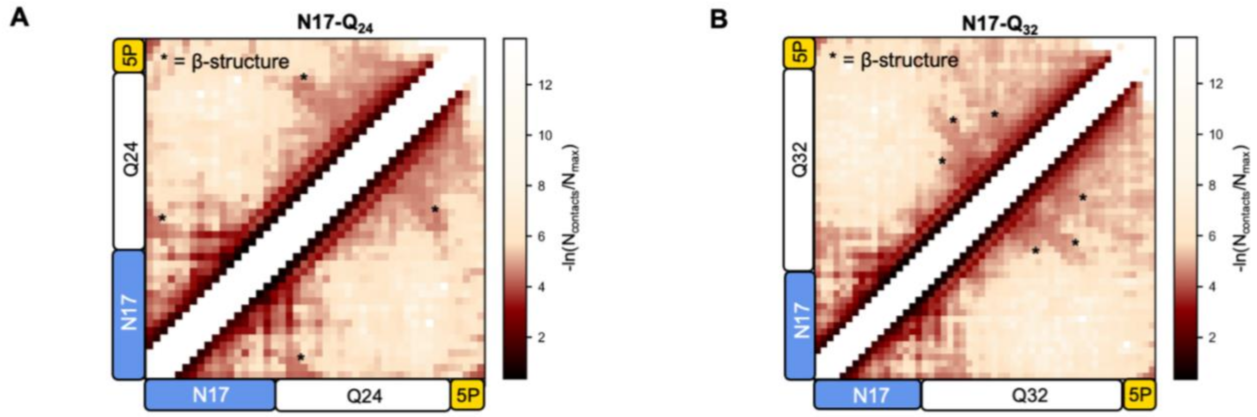


**Figure S7. Contact analysis of N17-polyQ$_{24/32}$ ensembles indicates transient β-sheet conformations.** Related to Figure 2.

**A**. Two-dimensional intramolecular contact maps calculated over three independent N17-polyQ$_{24}$ trajectories highlighting the transient (low) population of β-sheet conformations (marked as *) relative to $\alpha$-helices in the ensemble. **B.** Same as in A for N17-polyQ$_{32}$ trajectories.

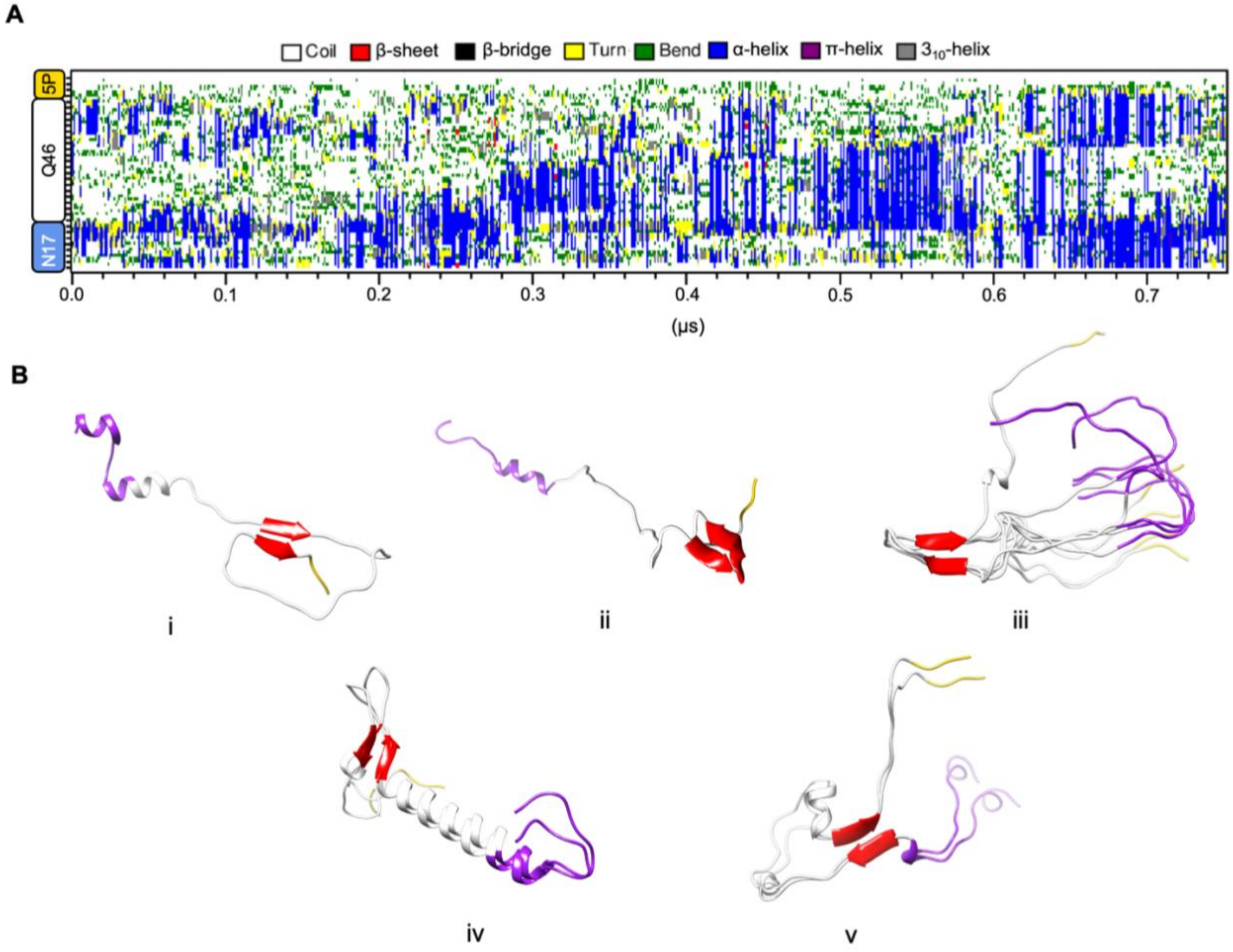


**Figure S8. Secondary structure analysis of the 293 K replica trajectory from PT-WTE simulations.** Related to Figure 2. **A.** Secondary structure variation as a function of time for N17-$Q_{46}$ from replica trajectory. **B**. Representative β-sheet structures or clusters (i-vi) which transiently form in the replica trajectory over 600 ns (first 150 ns excluded as equilibration time) analyzed every 100 ps. The coloring scheme is as described for Fig. S5C.

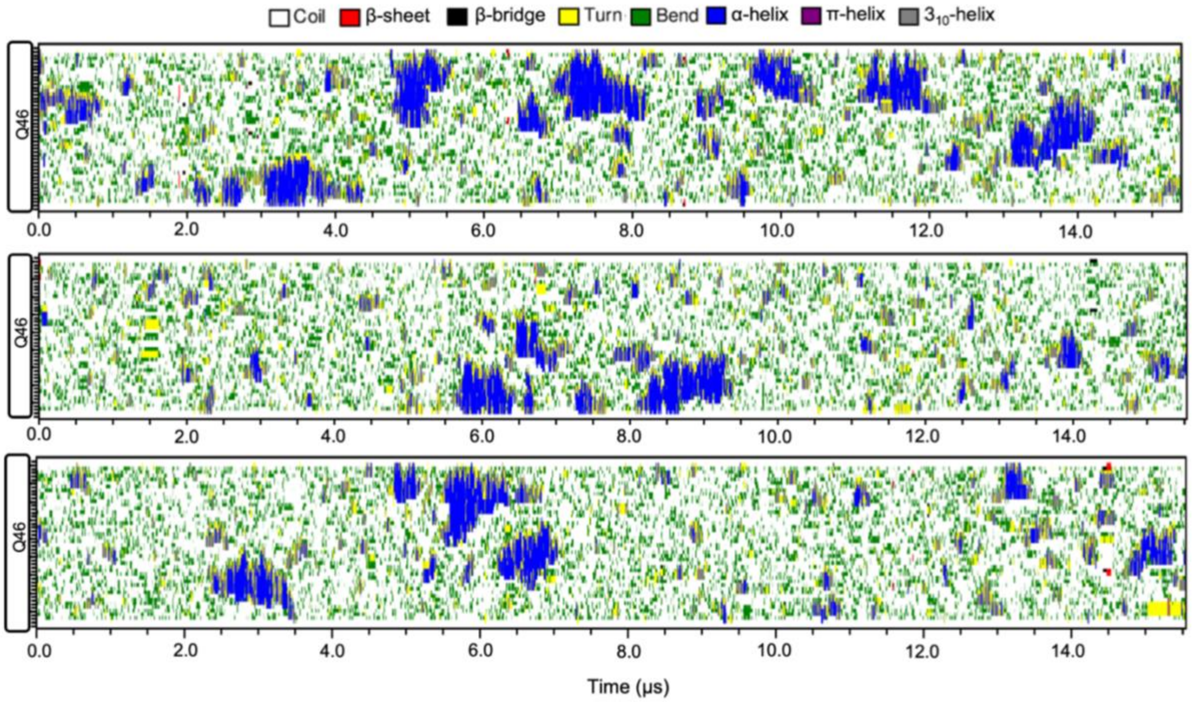


**Figure S9. Secondary structure variation in the $Q_{46}$ ensemble.** Related to Figure 2.
DSSP secondary structure variation as a function of time for $Q_{46}$ from three unbiased trajectories at 293.15 K.

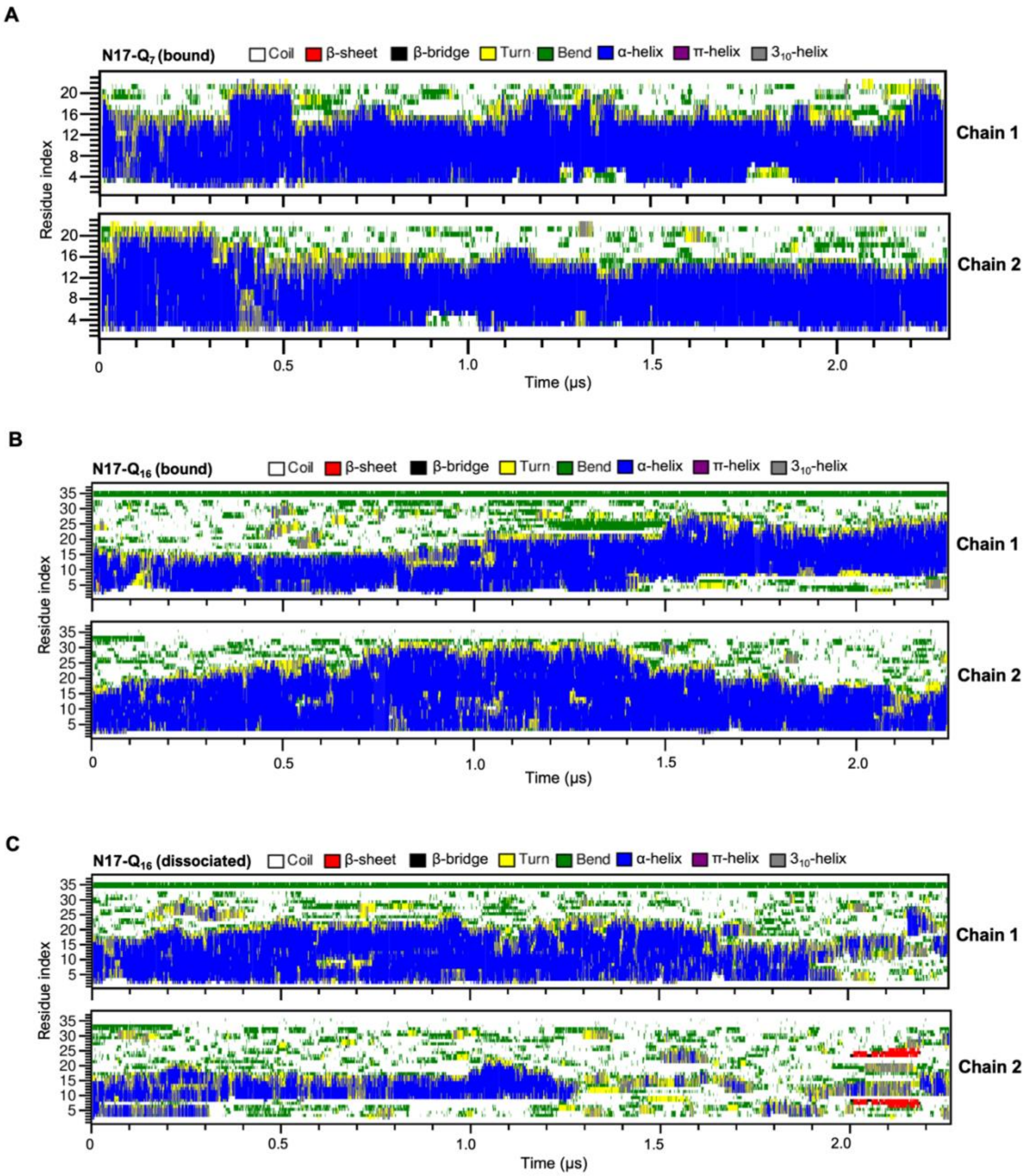


**Figure S10. Effect of dimerization on N17-$Q_{7/Q16}$ secondary structure stability.** Related to Figure 3

**A.** Secondary structure variation as a function of time for each chain in a representative N17-$Q_7$ dimer trajectory which shows stable association. The plot highlights the stabilization of α-helical conformations in the bound dimer. **B.** Same as in A for C for N17-$Q_{16}$. **C.** Secondary structure variation as a function of time for each chain in a representative N17-$Q_{16}$ dimer trajectory which undergoes dissociation after 1.3 μs. The plot highlights the destabilization of α-helical conformations upon dimer dissociation.

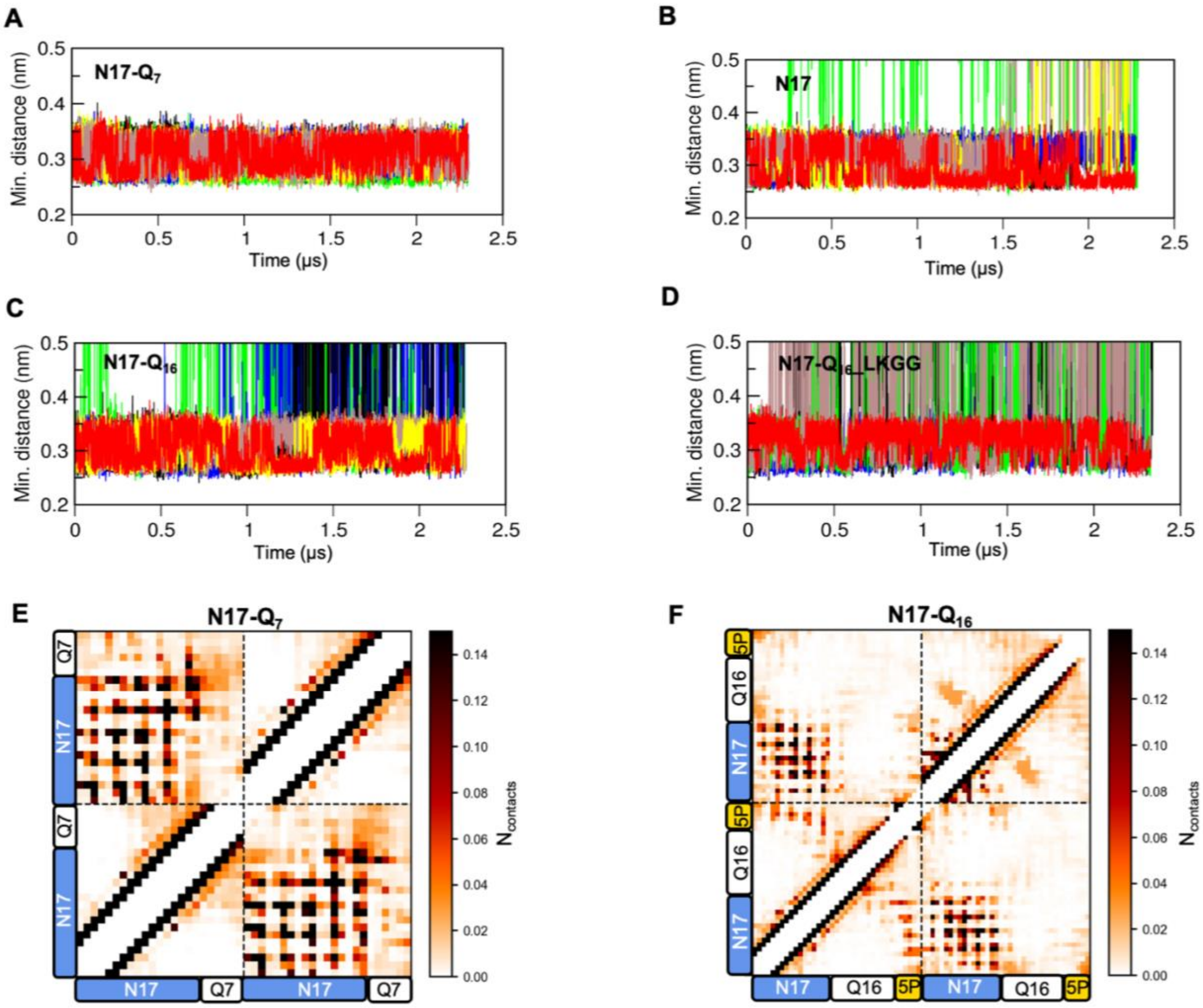


**Figure S11. Minimum distance and pairwise contact analysis to assess the effect of domain "cross-talk" on dimer stability.** Related to Figure 3

**A.** Minimum distance analysis as a function of time between monomer units of N17-$Q_7$ dimer. **B.** Same as in A for N17 dimer. Time points where the minimum distance first exceeds 0.4 nm corresponds to complete dissociation of the dimer. **C.** Same as in B for N17-$Q_{16}$. **D .** Same as in B for N17-$Q_{16}$_LKGG variant. **E**. 2D contact map highlight the network of weak inter-domain interactions within the N17-$Q_7$ dimer. The contact map was averaged over six independent trajectories. **F**. Same as in panel B for the N17-$Q_{16}$ dimer

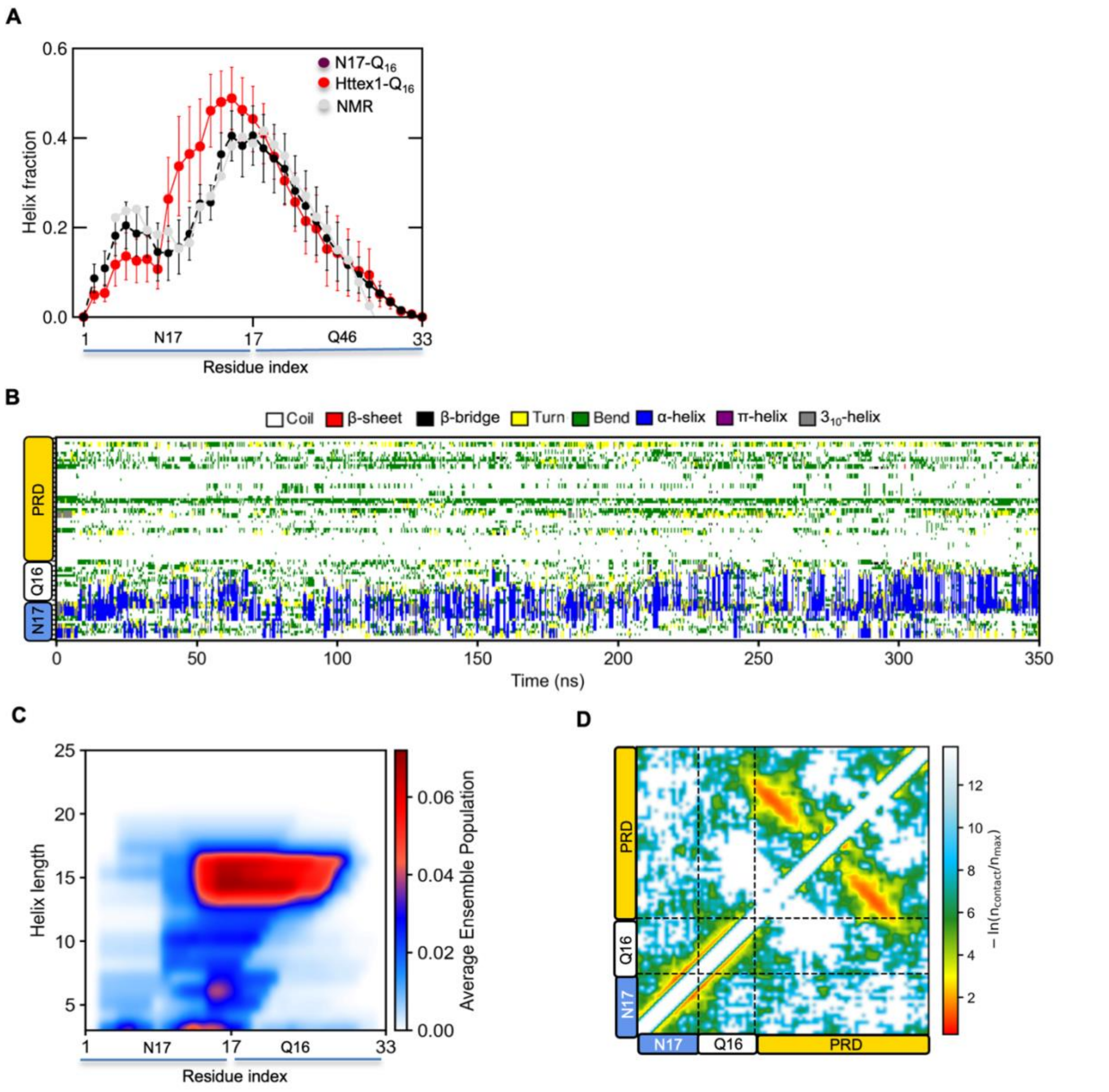


**Figure S12. Structure ensemble and intradomain interactions of Httex1-$Q_{16}$ computed from PT-WTE simulations.** Related to Figure 4.

**A**. Per-residue α-helical fraction of Httex1 (293 K replica) and its comparison with N17-$Q_{16}$ and NMR SSP scores. The first 100 ns was excluded as equilibration time. The error bars for Httex1 correspond to SEM over 4 independent blocks (62.5 ns each) of the 200 ns trajectory. **B**. DSSP secondary structure variation as a function of time for the 293 K replica trajectory. **C**. SS-map of Httex1-$Q_{16}$ (left) computed from the 293 K replica trajectory (excluding first 50 ns) indicating the probability of various α-helical lengths across N17 and polyQ regions. **D**. 2D-intramolecular contact map analysis of the Httex1 (293 K) ensemble highlighting the prevalence of PRD intradomain interactions involving two polyproline tracts ($P_{10}$/$P_{11}$).

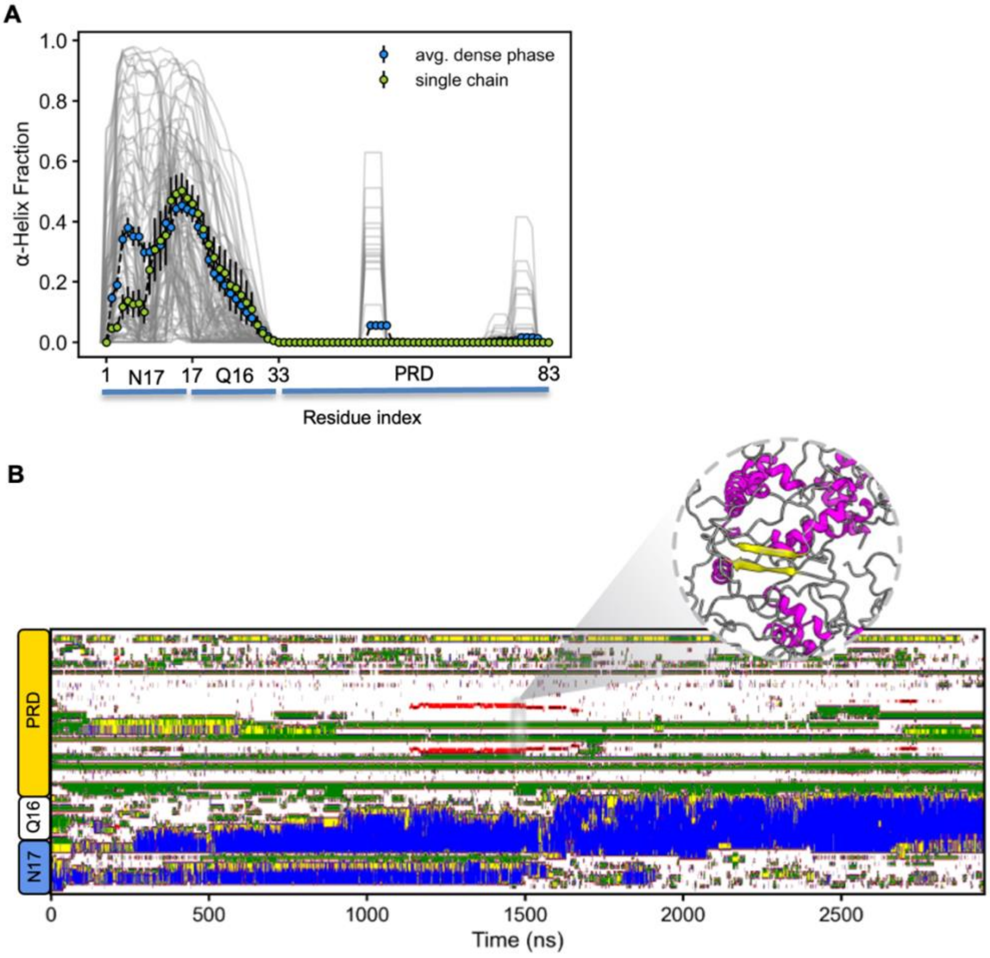


**Figure S13. Structural analysis of Httex1-$Q_{16}$ condensate.** Related to Figure 4.
**A**. Mean α-helical fraction per residue over all dense phase chains reveals a substantial enhancement in N17 α-helicity (aa: 2-11) compared to single chain (monomer). **B**. DSSP secondary structure variation as a function of time for a representative molecule within the condensate which forms a transient β-sheet from 1.2 to 1.5 us.
